# XtractPAV: An Automated Pipeline for Identifying Presence–Absence Variations Across Multiple Genomes

**DOI:** 10.1101/2025.06.27.661953

**Authors:** Rana Sheraz Ahmad, Muhammad Sadaqat, Muhammad Tahir ul Qamar

**Affiliations:** Integrative Omics and Molecular Modeling Laboratory, Department of Bioinformatics and Biotechnology, Government College University Faisalabad (GCUF), Faisalabad, 38000, Pakistan; UMR CNRS 6553 Ecosystèmes, Biodiversité, Evolution (ECOBIO), Université de Rennes 1, Rennes, France

## Abstract

**Motivation:** Presence-absence variations (PAVs) significantly influence phenotypic diversity across and within species by modulating functional modules involved in stress responsiveness, adaptation, and developmental processes. This modulation ultimately contributes to genetic diversity at both inter- and intra-species levels. However, existing tools for detecting PAVs offer limitations in achieving optimal analysis because they lack scalable workflows for multi-genome comparisons and frequently necessitate manual integration. To address these challenges, we developed XtractPAV, an end-to-end pipeline that automates the extraction, annotation, and interactive visualization of PAVs across large-scale genomic datasets.

**Results:** XtractPAV was evaluated using assembled genomes of both eukaryotic and prokaryotic organisms, including *Pyrus communis, Arabidopsis thaliana, Mus musculus*, and *Salmonella enterica*, to assess its ability to detect the genomic variations across diverse species. The performance of XtractPAV was benchmarked against other established pipelines, demonstrating superior precision and a more comprehensive extraction of PAV segments. Notably, our pipeline not only identified the known PAVs from the reference set but also revealed novel variations in genes associated with various functions such as flowering time regulation and disease resistance. Furthermore, the automated report generation feature of XtractPAV produces publication-ready summaries of PAV distributions and related metrics.

**Availability:** XtractPAV is freely accessible at https://github.com/SherazAhmadd/XtractPAV and on the XtractPAV webpage. The Package includes all requisite files, a user manual, test data, and a license permitting non-commercial use.

**Supplementary material:** Supplementary data are accessible online at *Bioinformatics*.

## 1. Introduction

Presence-absence variations (PAVs) represent a significant class of genomic structural variations (SVs), characterized by the presence of specific genomic segments in some individuals and their absence in others. These regions of variation are often associated with key functions or traits, thereby facilitating an understanding of phenotypic differences across species (Wang, et al., 2023). Thus, PAVs contribute to the sequence diversity among individuals, leading to intraspecific variability and interspecific genomic divergence (Gerdol, et al., 2025). The identification of large SVs is considerably more complex than that of small-scale genomic variations (Jiao, et al., 2025), often leading to their oversight in genomics studies. Various tools have been developed for the detection of these large-scale SVs, particularly PAVs, such as ScanPAV (Giordano, et al., 2018) and ppsPCP (Tahir Ul Qamar, et al., 2019). However, these tools impose strict limitations and are unable to provide comprehensive annotation of PAVs, including those intersecting coding regions. To address these limitations, this study introduces XtractPAV, a novel pipeline designed to efficiently and accurately trace PAVs. XtractPAV has been evaluated across five complex pear genomes, eighteen ecotypes of Arabidopsis, and six mouse genome assemblies to demonstrate its broad applicability and accuracy.

## 2. Materials and Methods

To identify the genomic regions present in the query genome but absent in the reference genome, a comprehensive whole-genome comparison is first performed. The NUCmer program from the MUMmer4 (Marçais, et al., 2018) is employed to compute the delta file of the alignment, which is subsequently analysed for coordinate information. In-house Python scripts are utilized to identify the unmapped regions in the genome, with the minimum length parameter adjusted by the user, and to retrieve the corresponding sequences. To address larger segments of ambiguous bases (N’s), PAVs are segmented into smaller fragments prior to the downstream filtering. To ensure the authenticity of the PAVs, all-versus-all alignment with the reference genome is performed using BLASTn (Camacho, et al., 2009); fragments with no hits, less than 10% coverage, and under 5% sequence identity are considered authentic PAVs and retained for further analysis. Those PAVs located within the coding regions are designated as the genic PAVs. Additional in-house Python scripts are employed to refine the boundaries of genic-PAVs and annotate them using available annotation file of the query genome. In XtractPAV, we extend the PAV sequence when it intersects with the coding region to complete the gene sequence, subsequently storing this information in a FASTA file. However, the coordinates in the summary file retain the original PAV designation. Ultimately, the pipeline produces a comprehensive final analysis report along with various interactive plotting graphs (Fig. 1). XtractPAV enables the simultaneous analysis of multiple query genomes against the reference genome in a single execution. While utilizing XtractPAV, users can specify parameters such as minimum length, coverage, similarity, thread count, and the number of genomes for PAV extraction.

**Fig. 1.**
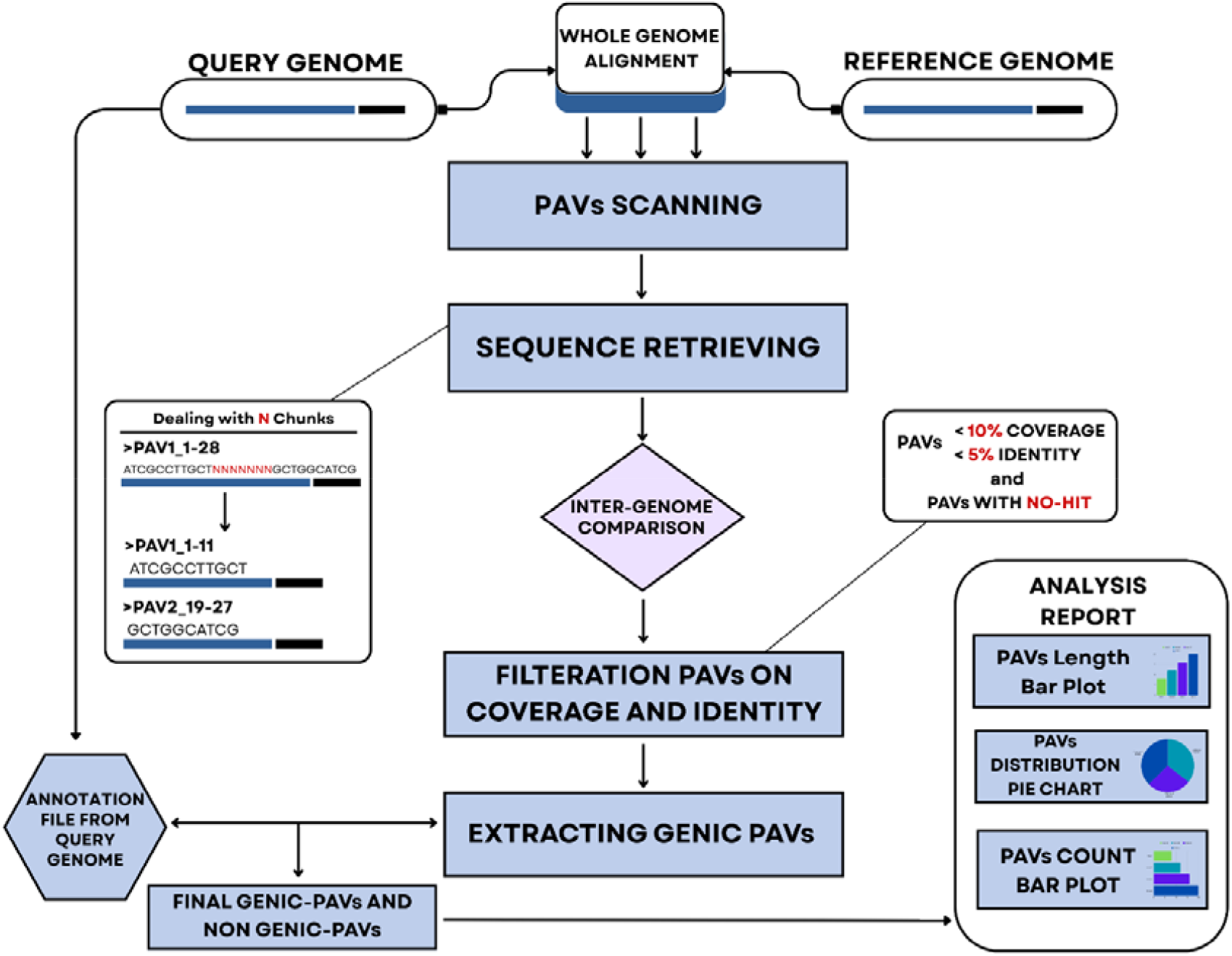
Workflow diagram of the XtractPAV pipeline. This pipeline accepts two genome assemblies in FASTA format, along with their corresponding annotation files in General Feature Format version 3 (GFF3), for the analysis of PAVs.

## 3. Results and Discussion

XtractPAV was developed to accommodate both eukaryotic and prokaryotic organisms based on user requirements. The scripts for XtractPAV are written in Python and Shell scripting and are implemented in a Linux environment. It accepts genomic sequences in *.fa, *.fna, *.faa, and other *.fasta formats, alongside their annotation file in GFF3, which is recommended; however, *.gff and *.gtf formats are also acceptable. All PAVs are recorded in FASTA format, accompanied by their annotations in GFF3 format and a local HTML file for analysis reports and graphical representations. The performance of XtractPAV was evaluated for both eukaryotic and prokaryotic organisms. For a complex plant genome analysis, five *Pyrus communis* (pear) genomes, with Dangshansuli serving as the reference, while Bartlett, Cuiguan, Shanxiduli, and ZhongaiNO.1 genomes, were sourced from PGDB (Chen, et al., 2023) to investigate interspecies differences, revealing a significant number of insertions and deletions among the species. Furthermore, we selected 19 ecotypes of Arabidopsis from the 19 genomes of the *Arabidopsis thaliana* project (Gan, et al., 2011) to conduct PAVs analysis, aiming to elucidate their genetic divergences and quantify the number of PAVs (Supplementary Table S1).

The mouse (*Mus musculus*) is a widely utilized model organism for the investigation of human diseases and biological processes; however, the presence of multiple genome assemblies and substantial intraspecies genetic variation contributes to phenotypic diversity, which may influence empirical findings. To investigate SVs across various mouse assemblies, we conducted the PAVs analysis using XtractPAV on six mouse assemblies, AKR_J, 129S1_SvImJ, C3H_HeJ, C57BL_6NJ, and DBA_2J, compared to the reference genome GRCm39 (*mm39*). To test the performance of XtractPAV on prokaryotic genomes, we selected 42 serovars of *Salmonella enterica*, which were previously employed to construct the pan-genome (Jacobsen, et al., 2011). *S. enterica typhimurium* strain LT2 served as the reference, while the other 41 strains were utilized as target genomes. The results of the *S. enterica* PAV analysis are presented in Supplementary Table S2.

### 3.1 Benchmarking

To assess the functional capabilities of XtractPAV, we conducted a comparative analysis with other PAV extraction pipelines, specifically ScanPAV and ppsPCP. While these tools can be effective in certain contexts, they also demonstrate limitations in managing multi-genomic data, annotating coding-regions, adjusting parameter flexibility, and accurately identifying PAVs. All tools were evaluated using the same dataset of *P. communis* (Pear) genomes to ensure consistent processing (Table 1). The discrepancies in results when comparing different pipelines can be attributed to the differences in their stringent identification and filtering criteria.

**Table 1.** Number of presence-absence variations (PAVs) identified with different pipelines to assess the performance of XtractPAV.

| Assembly | Raw PAVs |  |  | Validated PAVs |  |  |
| --- | --- | --- | --- | --- | --- | --- |
|  | XtractPAV | ppsPCP | ScanPAV | XtractPAV | ppsPCP | ScanPAV |
| *Bartlett* | 43,699 | 43,235 | 36,794 | 24,377 | 23,291 | 36,794 |
| *Cuiguan* | 32,262 | 31,972 | 25,304 | 13,207 | 18,527 | 25,304 |
| *Shanxiduli* | 38,210 | 37,855 | 28,898 | 16,168 | 21,822 | 28,898 |
| *ZhongaiNO.1* | 37,134 | 36,806 | 28,870 | 16,720 | 19,955 | 28,870 |

The scanPAV pipeline is specifically designed for the extraction of PAVs. However, it presents several limitations. Notably, it cannot process multi-genomic inputs, and the size of the PAV is fixed at 1 kilobase (kb). While PAVs in plants have been documented to be a minimum of 100 base pairs (bps) (Shen, et al., 2015). Since users cannot adjust the length of PAVs according to the specific genome and analysis requirements, which renders it is predominantly suitable for the mammalian genomes. Furthermore, scanPAV does not explicitly extract coding regions along with their annotations. Additionally, the pipeline has not incorporated validation based on coverage and similarity, which raises concerns regarding its precision.

XtractPAV represents an advanced iteration of our previous pan-genome construction pipeline, ppsPCP (Tahir Ul Qamar, et al., 2019), and is specifically designed to identify the PAVs with improved features and performance. The ppsPCP tool was initially developed for constructing the pangenomes, but it is often utilized to screen PAVs (Chen, et al., 2025; Lan, et al., 2024). However, it imposes a strict minimum length of 100 bp for PAVs, which cannot be modified by the user depending on specific genome and analysis needs. Additionally, since ppsPCP is primarily tailored for pangenome construction, which contributes to longer processing times for PAV extraction and utilizes the primary PAV output as an intermediary file, it does not provide PAVs as explicit output and annotation of genic PAVs. In contrast, XtractPAV addresses these limitations by offering a more comprehensive solution for PAV extraction. It accepts multi-genomic input, allows for fully customizable parameters, and provides direct outputs including PAVs in FASTA format, accompanied by an annotation GFF file, and a graphical report that summarizes the PAVs distribution across genomes.

## 5. Conclusion

The consideration of complete diversity and genetic composition, without accounting for PAV regions, presents an insufficient representation of an organism’s genetic architecture. The challenge of identifying large SVs remains significant. XtractPAV addresses this issue by facilitating the tracking the larger-scale insertions and deletions, particularly in PAV regions, across multiple genomes, utilizing customizable user-defined parameters. Furthermore, by quantifying both genic and non-genic PAVs, XtractPAV also offers a robust metric to evaluate genome assembly quality across various sequencing technologies.

## Supporting information

analysis are presented in Supplementary Table 2.

## Acknowledgment

Not applicable.

## Author contributions

R.S.A.: logical programming, coding, code testing, visualization, and writing the manuscript. M.S.: code testing, validation, and editing of the manuscript. M.T.Q.: conception and study design, code testing, resources, supervision, and editing the manuscript.

## Supplementary data

Supplementary data are available at Bioinformatics online.

## Ethical Statement

This study did not involve any experiments with animal subjects or human participants. The authors utilized ChatGPT v2 to enhance the clarity and readability of the codes. All content generated through this service was carefully reviewed and edited by the authors, who take full responsibility for the final content of the publication.

