## Supplementary material for "XtractPAV: An Automated Pipeline for Identifying Presence–Absence Variations Across Multiple Genomes": analysis are presented in Supplementary Table 2.

**1. Pipeline Implementation and Parameters**

1.1. Required Dependencies

 – MUMmer4

– BLAST+
 – Bedtool v2.27.1

 – Biopython 1.78

 – Plotly 6.0.1
 – Hardware specifications are subject to the input file data size and the computational power required for processing.

1.2. Default Flags

| **FLAG** | **Description** | **Default** |
| --- | --- | --- |
| --rf | Reference genome FASTA file | Required |
| --ra | Reference GFF3 annotation | Required |
| --qf | Query genome FASTA files (For Multi Genomes use comma‑sep) | Required |
| --qa | Query GFF3 annotation files (For Multi Genomes use comma‑sep) | Required |
| --cov | Minimum coverage threshold (0–1) | 0.8 ( Float ) |
| --sim | Minimum similarity percentage | 90.0 ( Float ) |
| --len | Minimum PAV length (bp) | 100 (integer) |
| --thr | Number of threads | 1 |
| --help | Show help message | Optional |
| --version | Show pipeline version | Optional |

1.3. Usage

XtractPAV.sh –rf reference_genome.fa --ra reference_annotation.gff3 –qf query1.fa, query2.fa –qa query1.gff3,query2.gff3 --cov 0.8 --sim 90.0 --len 100 --thr 8

**2. PAVs Extraction Criteria and genomes information**

2.1 *Arabidopsis Thaliana* Ecotypes

*A. Thaliana* became a model organism to study Plant Biology. It's straight molecular genetics that enabled prominent discoveries in plant development (Meinke, et al., 1998). We collected the eighteen ecotypes of *A. thaliana*, which are from different places such as Italy, Germany, Scotland, and Russia. etc. *A. thaliana* Whole-Genome Sequencing assemblies obtained from <http://mtweb.cs.ucl.ac.uk/mus/www/19genomes/>, which is 19 genomes of the Arabidopsis thaliana project (Gan, et al., 2011) in FASTA format along with their GFF3 annotation file. We choosed the Col-0 (Canary Isles ) as reference genome and Bur-0, Can-0, Ct-1, Edi-0, Hi-0, Kn-0, Ler-0, Mt-0, No-0, Oy-0, Po-0, Rsch-4, Sf-2, Tsu-0, Wil-2, Ws 0, Wu-0 and Zu-0 as the query genomes. We have run the XtractPAV on 24 threads and 100G memory with 90% coverage, 95% similarity, and 100 bps minimum length. It took just 53 minutes (0.89 CPU hours) to identify PAVs on all Arabidopsis ecotypes. The size of genomes is shown in Table 2.1.

**Table 2.1.** Arabidopsis thaliana Ecotype Accessions: Genome Assembly Details and Size

| **Accession** | **Version** | **Genome Size** |
| --- | --- | --- |
| Col_0 | v7 | 115 MB |
| bur_0 | v7 | 114 MB |
| can_0 | v7 | 113 MB |
| ct_1 | v7 | 114 MB |
| edi_0 | v7 | 115 MB |
| hi_0 | v7 | 114 MB |
| kn_0 | v7 | 114 MB |
| ler_0 | v7 | 114 MB |
| mt_0 | v7 | 113 MB |
| no_0 | v7 | 114 MB |
| oy_0 | v7 | 113 MB |
| po_0 | v7 | 114 MB |
| rsch_4 | v7 | 114 MB |
| sf_2 | v7 | 114 MB |
| tsu_0 | v7 | 114 MB |
| wil_2 | v7 | 113 MB |
| ws_0 | v7 | 114 MB |
| wu_0 | v7 | 114 MB |
| zu_0 | v7 | 114 MB |

2.2 *Pyrus communis* (Pear)

Pear belongs to the Rosaceae family, with the most significant temperate fruit worldwide. It has five major domesticated cultivars for production: P. communis, P. pyrifolia, P. bretschneideri, P. ussuriensis, and P. ussuriensis × P. communis. We selected all these for PAV analysis to investigate their diversity. All genomes were retrieved from the <http://pyrusgdb.sdau.edu.cn/>, which is PGDB (Chen, et al., 2023) in FASTA format with their annotation, respectively. We chose P. bretschneideri as the reference genome and P. communis, P. pyrifolia, P. ussuriensis, and P. ussuriensis × P. communis as query genomes, and XtractPAV was run on a cluster with the same parameters as Arabidopsis, it took 15 CPU hours to finish the analysis. We have also executed the job without the multi-genome option, and on average, each genome took 5.15 CPU hours; this shows that utilizing the multi-genome feature is more time-saving. Rest of the genome information is given below in Table 2.2

**Table 2.2.** *P. communis* (Pear) Accessions: Genome Assemblies and Detailed Information

| **Accession** | **Specie** | **Version** | **Genome size** |
| --- | --- | --- | --- |
| Dangshansuli ( Reference ) | *Pyrus bretschneideri* | v1.1 | 493 MB |
| Bartlett | *Pyrus communis* | V2.0 | 475 MB |
| Cuiguan | *Pyrus pyrifolia* | v1.0 | 521 MB |
| Shanxiduli | *Pyrus betulifolia* | v2.0 | 513 MB |
| ZhongaiNO.1 | *P. ussuriensis × P. communis* | v1.0 | 491 MB |

2.3 *Mus musculus* (Mouse)

 The mouse has been one of the prominent model organisms in genetics to study mammalian biology and understand the disease mechanism (Guénet, 2005). This extensive resource is now available for the collaborative cross and high-throughput mutation projects and makes the model organism suitable for studying the genetic variation, including PAVs. We performed a PAV analysis on the mouse organism with its different assemblies. GRCm39 was selected as the reference genome, and the 129S1_SvImJ_v1, AKR_J_v1, C3H_HeJ_v1, C57BL_6NJ_v1, and DBA_2J_v1 as the query genomes to process. XtractPAV runs with 24 threads, and 500 GB of memory was allocated to this task. For the mammal genome, we increased the minimum length of PAV to 150bp, 90% coverage, and 95% similarity criteria were enforced. All mouse genomes were collected from the Ensembl database ( <https://www.ensembl.org/info/about/species.html> ). XtractPAV took 32.6 CPU hours to identify PAVs across all accessions. The assembly's details are given in Table 2.3

**Table 2.3.** Overview of *M. musculus* (Mouse) Accessions, Genome Assemblies, and Key Characteristics

| **Accession** | **Assembly** | **Version** | **Genome size** |
| --- | --- | --- | --- |
| GCA_000001635.9 | GRCm39 | v1 | 2.58 GB |
| GCA_921998555.2 | 129S1_SvImJ | v1 | 2.58 GB |
| GCA_922000895.2 | AKR_J | v1 | 2.56 GB |
| GCA_921997125.2 | C3H_HeJ | v1 | 2.55 GB |
| GCA_921999865.2 | C57BL_6NJ | v1 | 2.65 GB |
| GCA_921998315.2 | DBA_2J | v1 | 2.46 GB |

**3. Supplementary Results** 3.1 Arabidopsis Ecotype PAV Summary

**Ecotype_1_ bur_0**
Total PAVs: 1093
Longest PAV: 2.7 kb

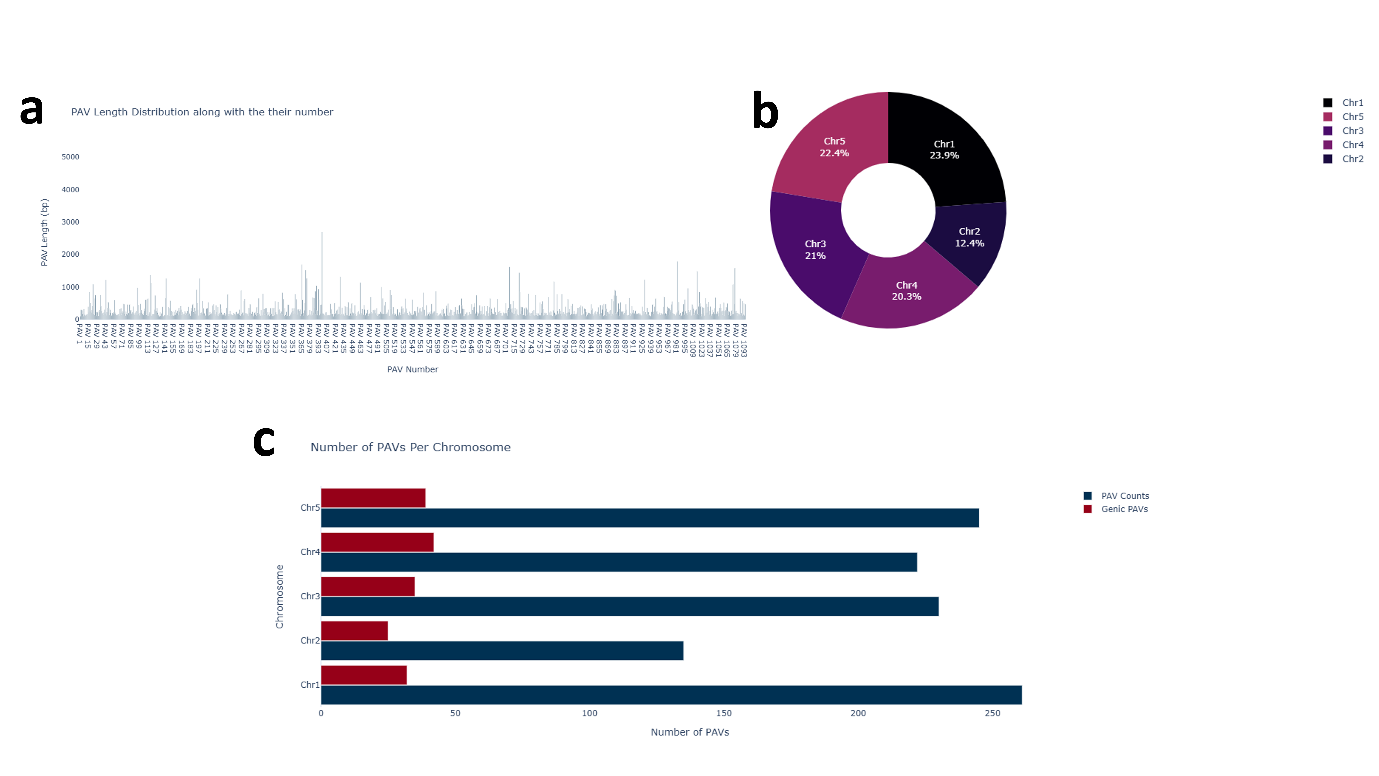

**Fig 3.1.** Result figures for the bur_0 assembly **(a)** Shows the number of PAVs with their length, highlighting both the shortest and longest detected regions, PAV number 398 is longest and it corresponds to chromosomal region **(b)** Percentage area covered by PAV on chromosomes **(c)** Classification of PAVs into genic and non-genic PAVs categories with their respective counts and 261 PAVs detected on chr1 on bur_0 assembly

**Ecotype_2_can_0**
Total PAVs: 1370
Longest PAV: 3.7 kb

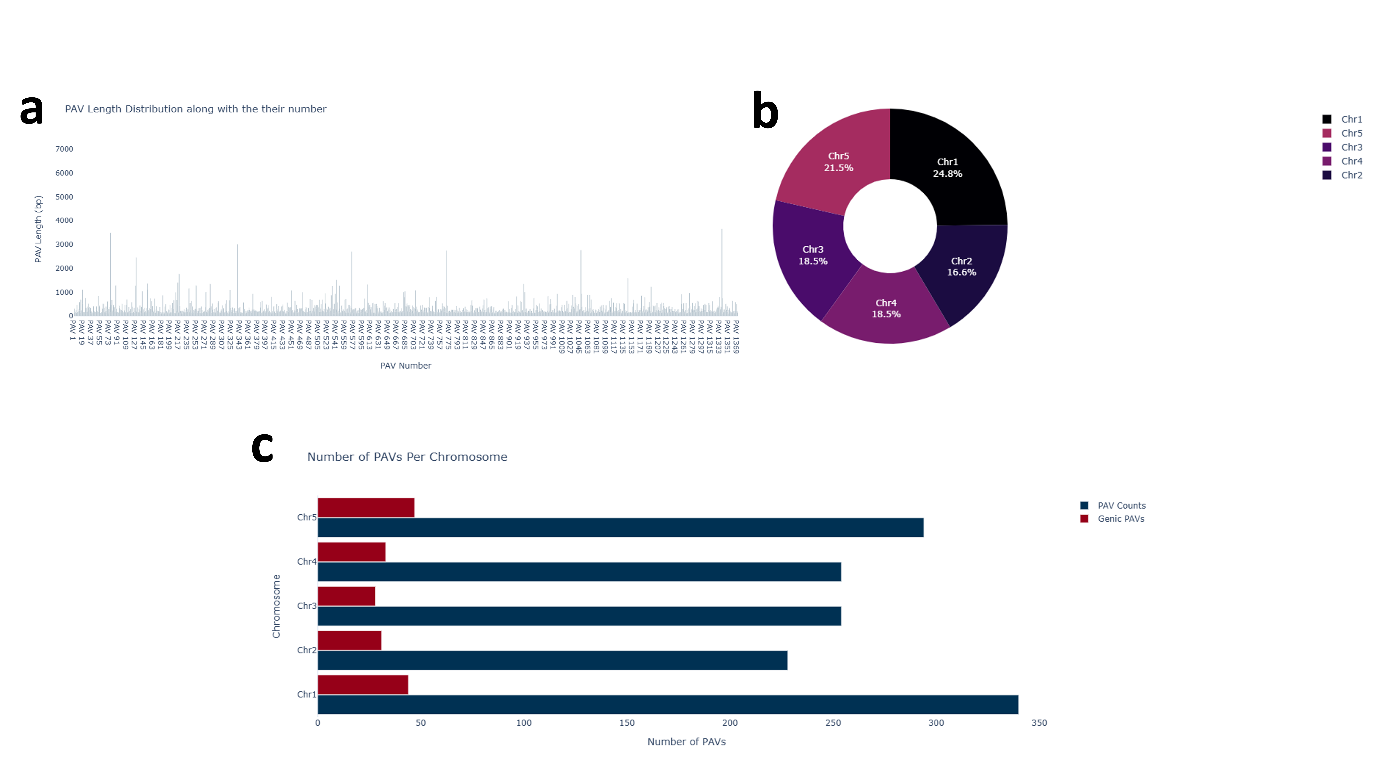

**Fig 3.2.** Summary of PAV analysis in the can_0 assembly **(a)** represents the number of Presence-Absence Variants (PAVs) by their length, highlighting both the shortest and longest detected regions, 1338 number PAV with 3362 bp found the longest **(b)** proportion of chromosomal area covered by PAV **(c)** Classification of PAVs into genic and non-genic categories with their respective counts.

**Ecotype_3_ct_1**
Total PAVs: 611
Longest PAV: 2.7 kb

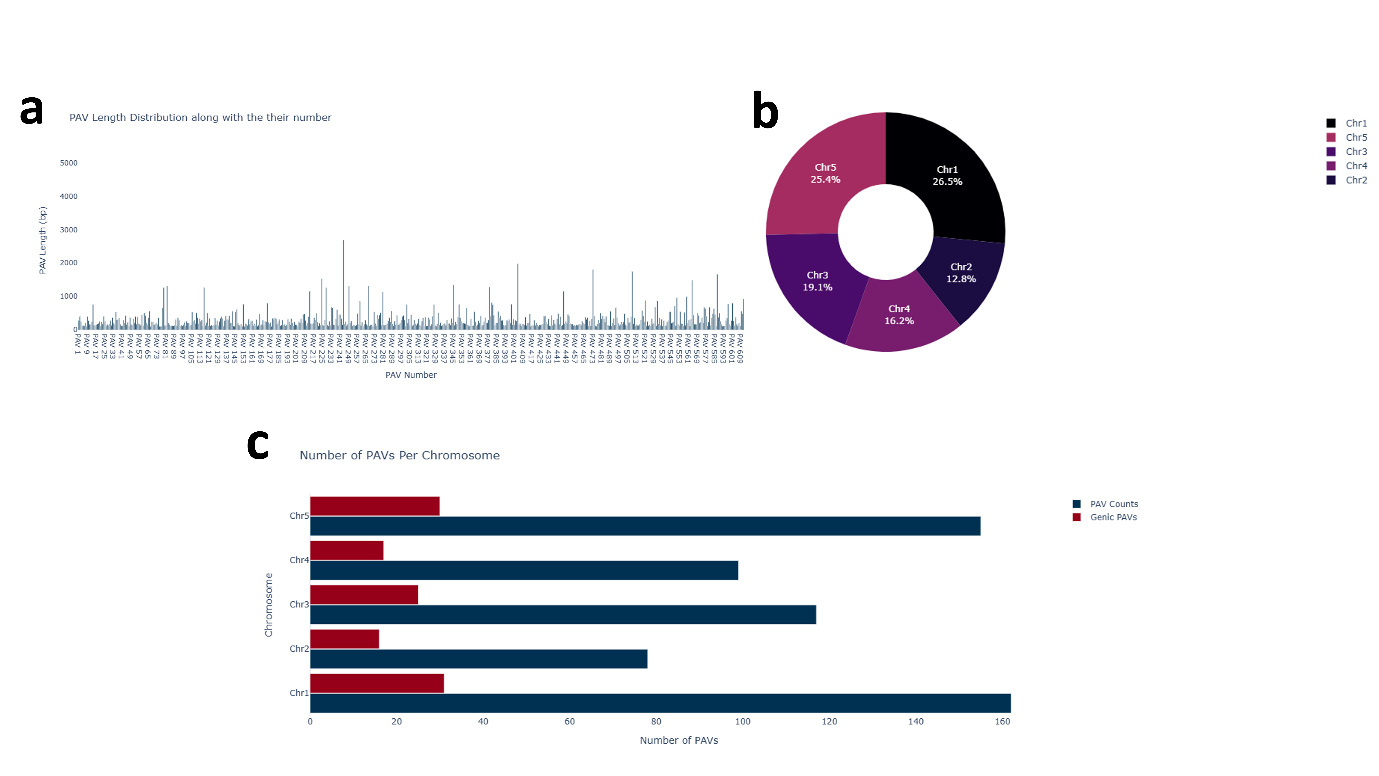

**Fig 3.3.** Overview of PAV characteristics in the ct_1 assembly **(a)** Length distribution of PAVs, PAV-244 is the longest gap region on chr3 **(b)** Chromosome number 1 has the highest number of missing regions as compared to other chromosomes **(c)** 31 genic PAVs have been found on chromosome number 1 in the ct_1 assembly.

**Ecotype_4_** **edi_0**
Total PAVs: 1201
Longest PAV: 6.1 kb

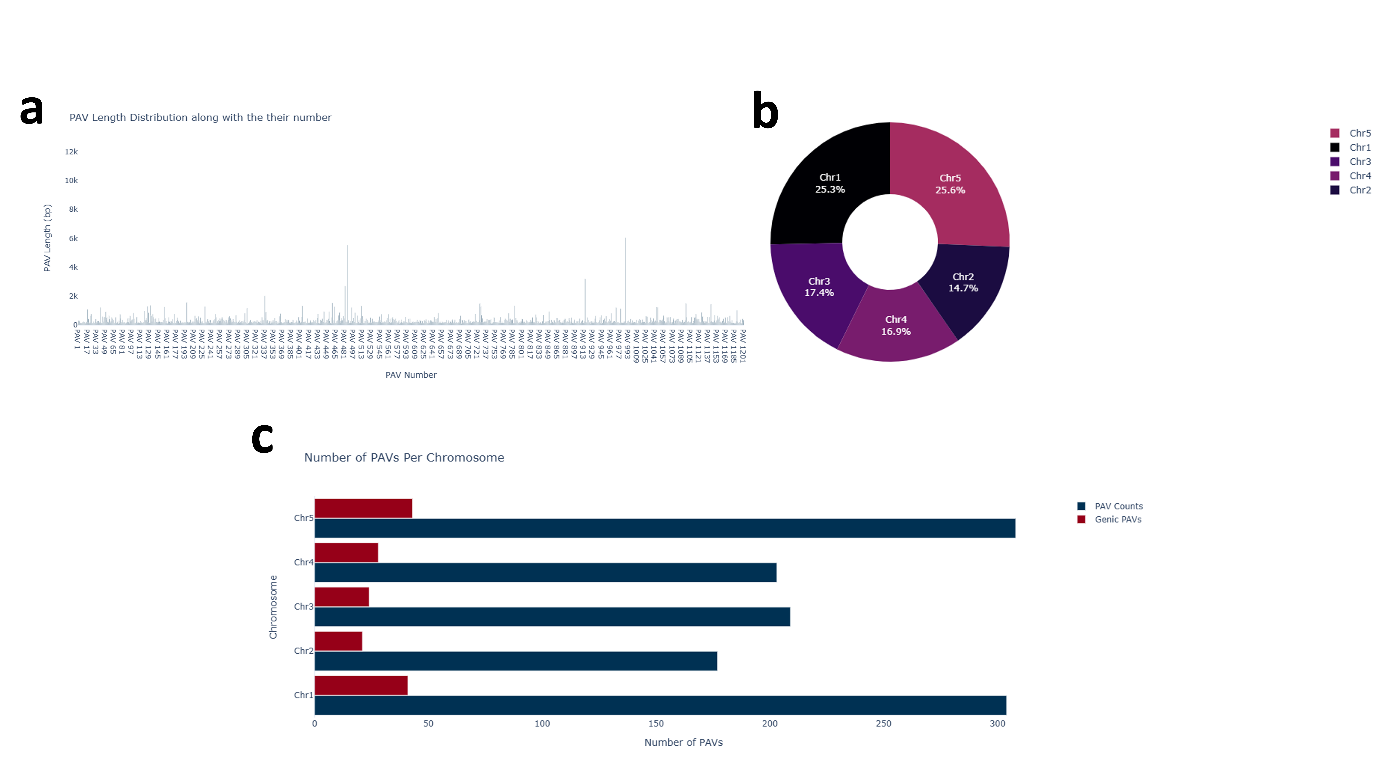

**Fig 3.4.** Figures showing the results for the edi_0 assembly **(a)** This figure reveal number of PAVs along with their length, PAV-988 is the longest and located on chromosome 1**(b)** Percentage area covered by PAVs on chromosomes **(c)** Classification of PAVs into genic and non-genic categories with their number of PAVs for each chromosome.

**Ecotype_5_hi_0**
Total PAVs: 450
Longest PAV: 2.7 kb

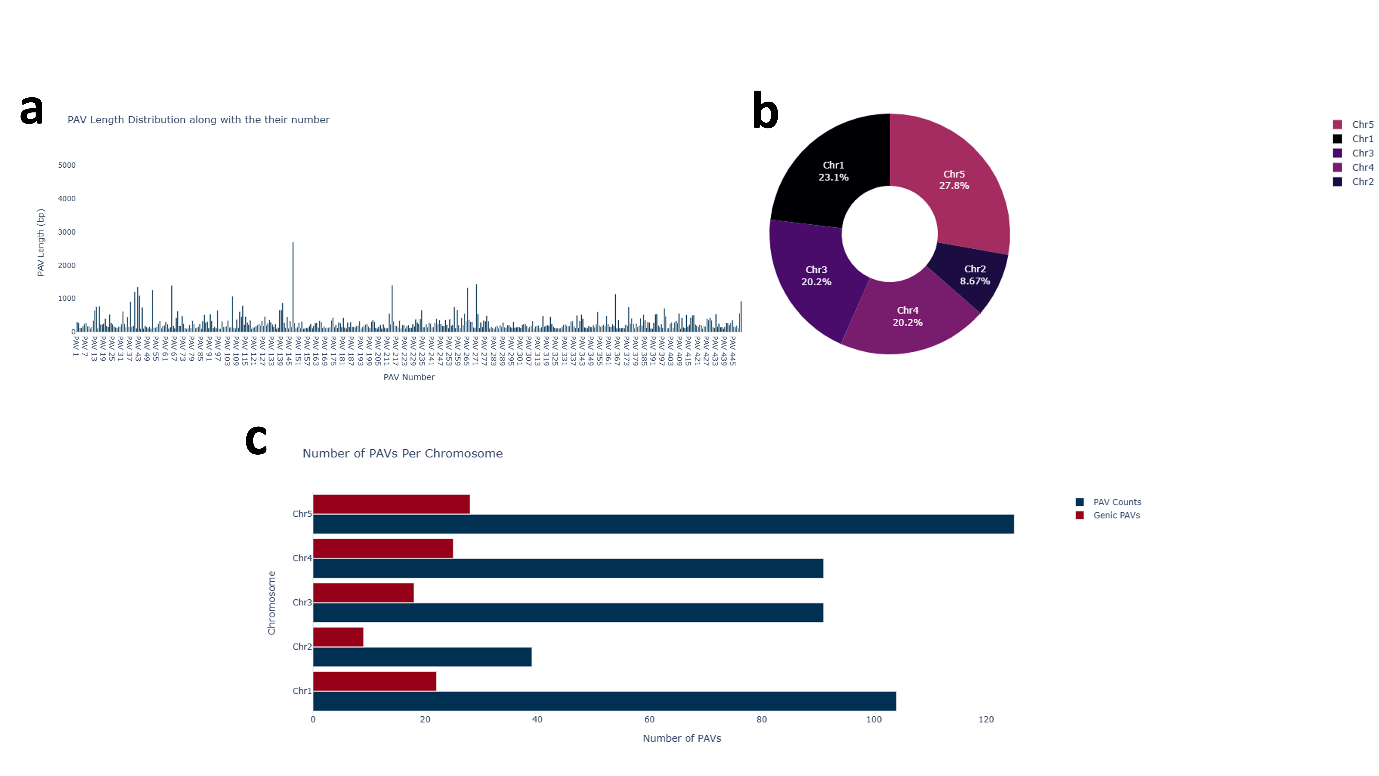

**Fig 3.5.** Characterization of PAVs in hi_0 assembly **(a)** shows the number of PAVs with their length, PAV number 147 has detected the longest missing (chromosomal) segment **(b),** proportion of chromosome occupied by PAV **(c).** Differentiating PAVs into genic and non-genic PAVs with counts provided.

**Ecotype_6_ kn_0**
Total PAVs: 1028
Longest PAV: 2.7 kb

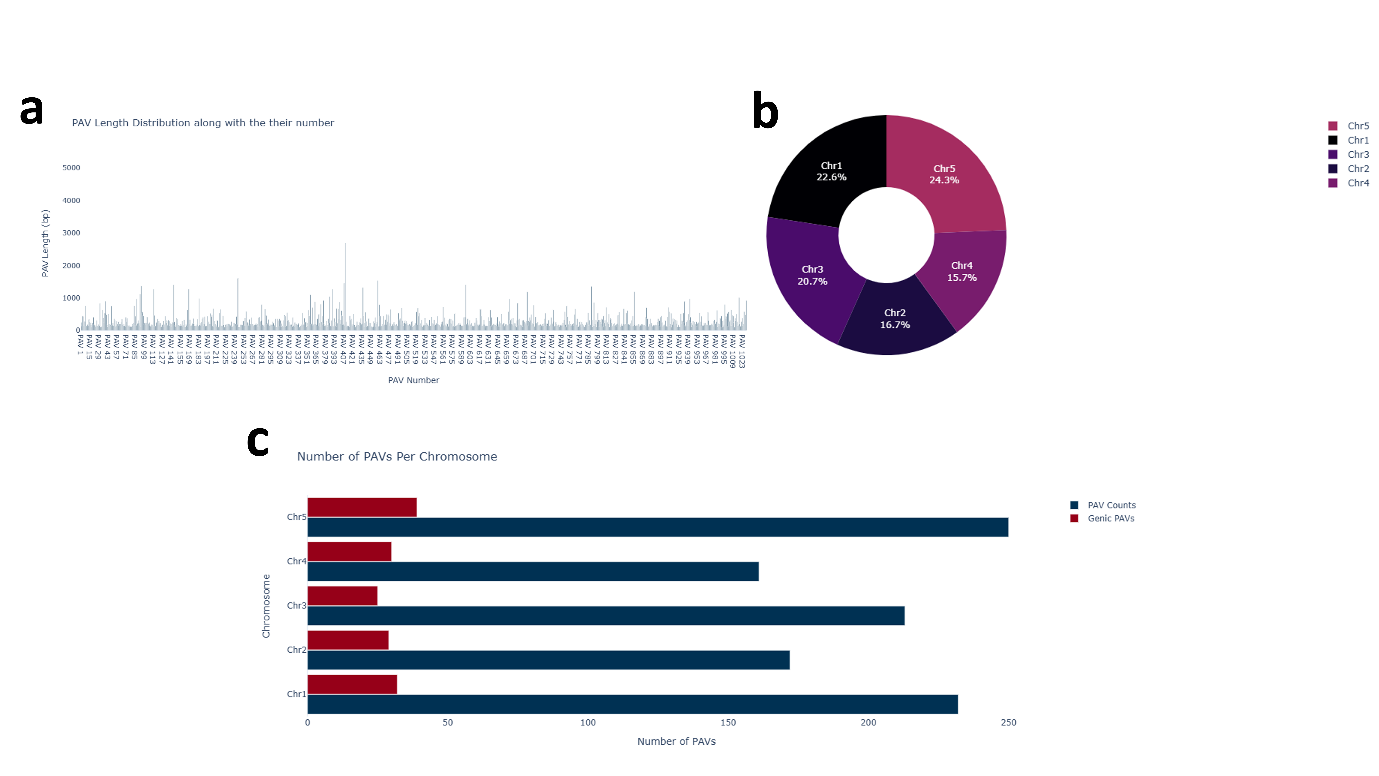

**Fig 3.6.** Analysis of Presence/Absence Variants (PAVs) in the kn_0 assembly **(a)** Quantifies the PAVs by length, PAV number 409 identified as the longest chunk **(b)** Shows the percentage of chromosomal region covered by PAVs **(c)** categorized PAVs into genicPAVs and non-genicPAVs type, providing the counts and notably, the chromosome 5 exhibits the highest level of coding variation in kn_0 assembly.

**Ecotype_7_ler_0**
Total PAVs: 1015
Longest PAV: 5 kb

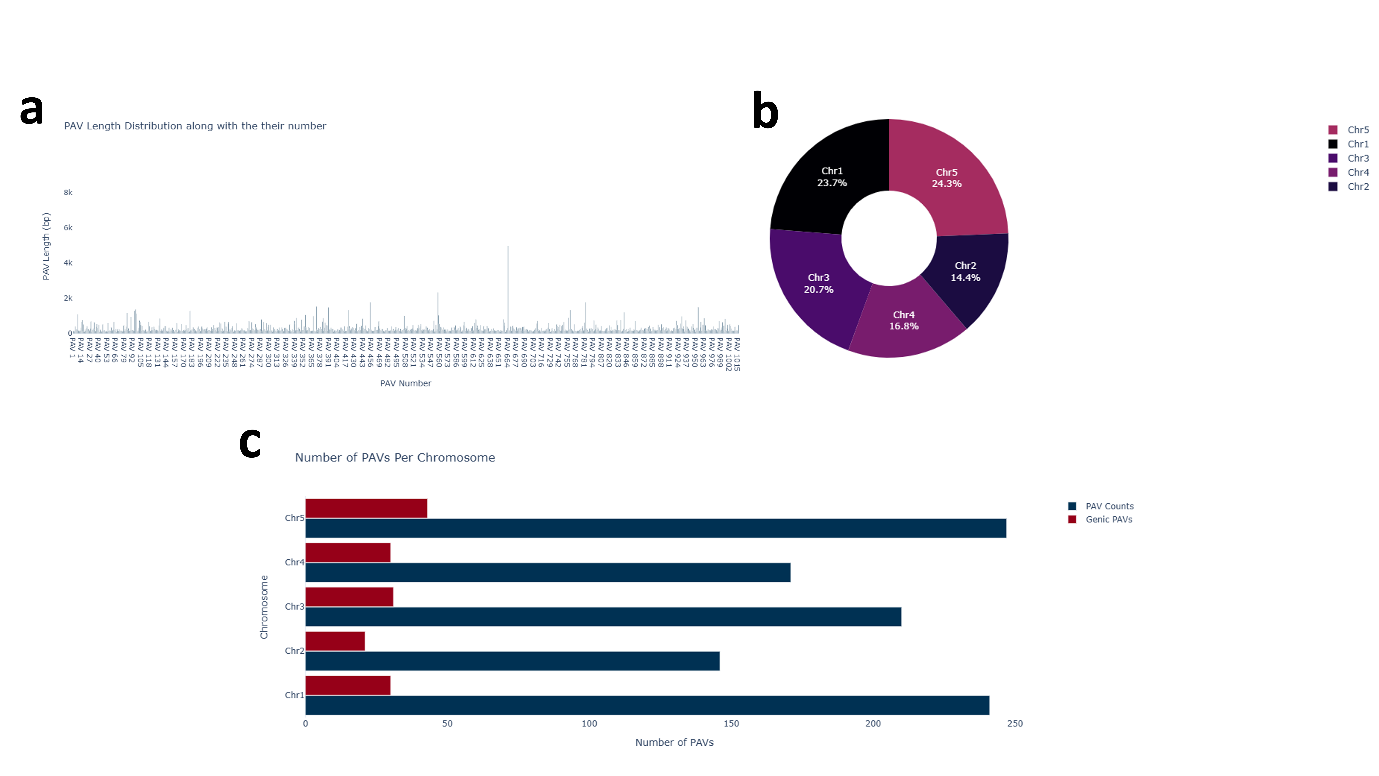

**Fig 3.7.** Provides an overview of presence-absence variants in the ler_0 assembly **(a)** presents PAV counts by length, the PAV number 664 is longest **(b)** Shows the percentage of chromosome coverage by PAVs **(c)** Classifies PAVs into genic and non-genic PAVs categories and the large number of PAVs are found on chromosome 1, 3, and 5 in ler_0 assembly.

**Ecotype_8_mt_0**
Total PAVs: 543
Longest PAV: 1.7 kb

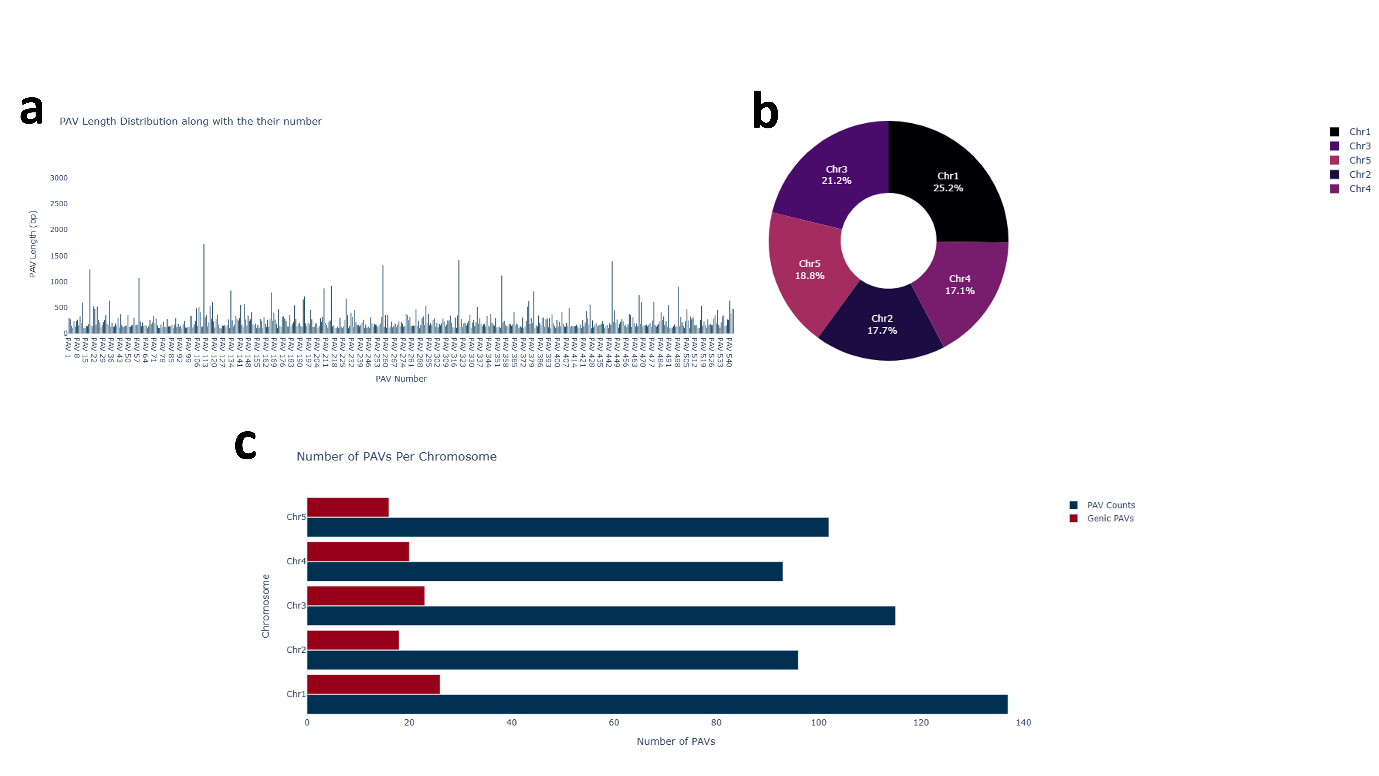

**Fig 3.8.** Result figures for the mt_0 assembly **(a)** show the number of PAVs with their length, PAV number 111 identified as the longest segment that is missing in the reference **(b).** The percentage area covered by PAV on chromosomes **(c)** divides the PAVs into genic and non-genic PAVs, corresponding to their counts.

**Ecotype_9_no_0**
Total PAVs: 1038
Longest PAV: 3.5 kb

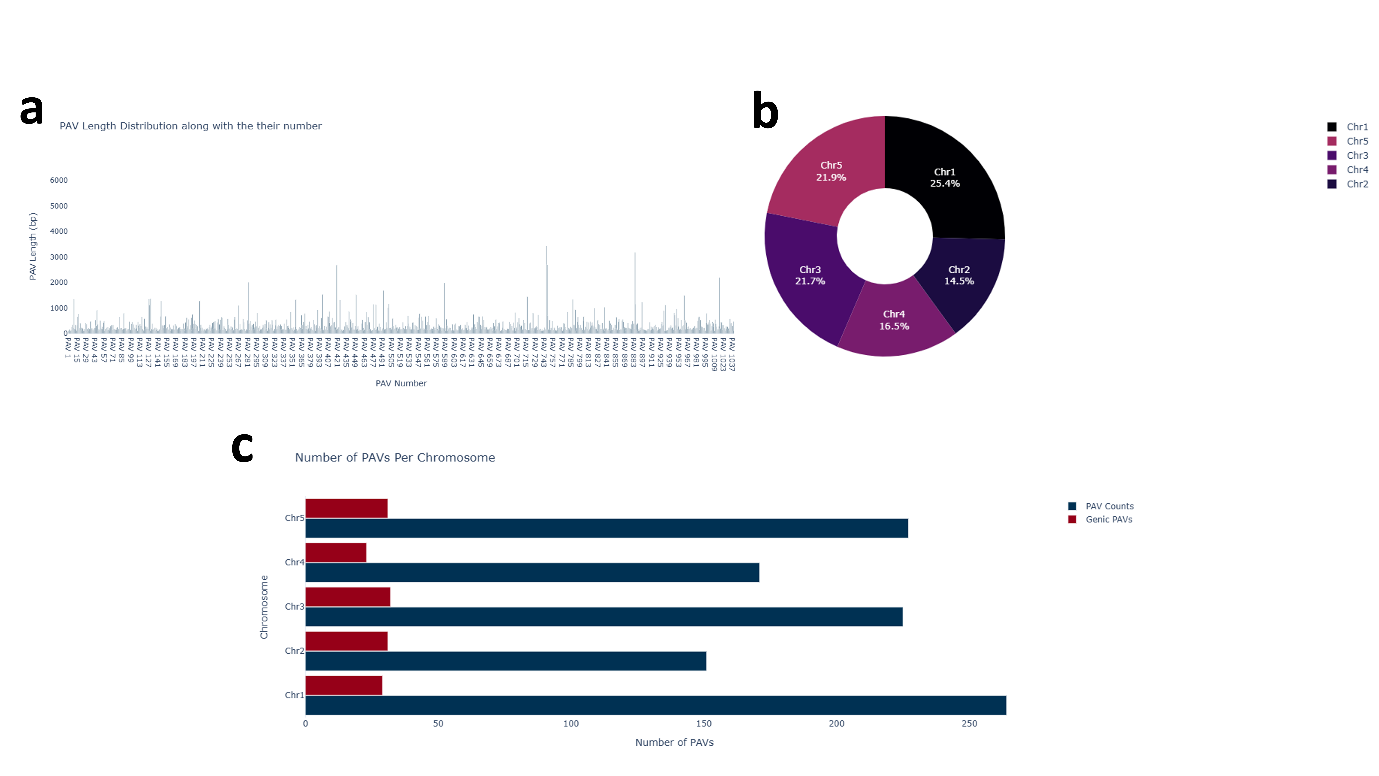

**Fig 3.9.** Result figures for the no_0 assembly **(a)** Shows the number of PAVs with their length, highlighting both the shortest and longest detected regions, PAV number 749 is longest **(b)** Percentage area covered by PAV on chromosomes **(c)** PAVs were assigned to genic and non-genic categories, and the respective counts were recorded.

**Ecotype_10_oy_0**
Total PAVs: 564
Longest PAV: 2.7 kb

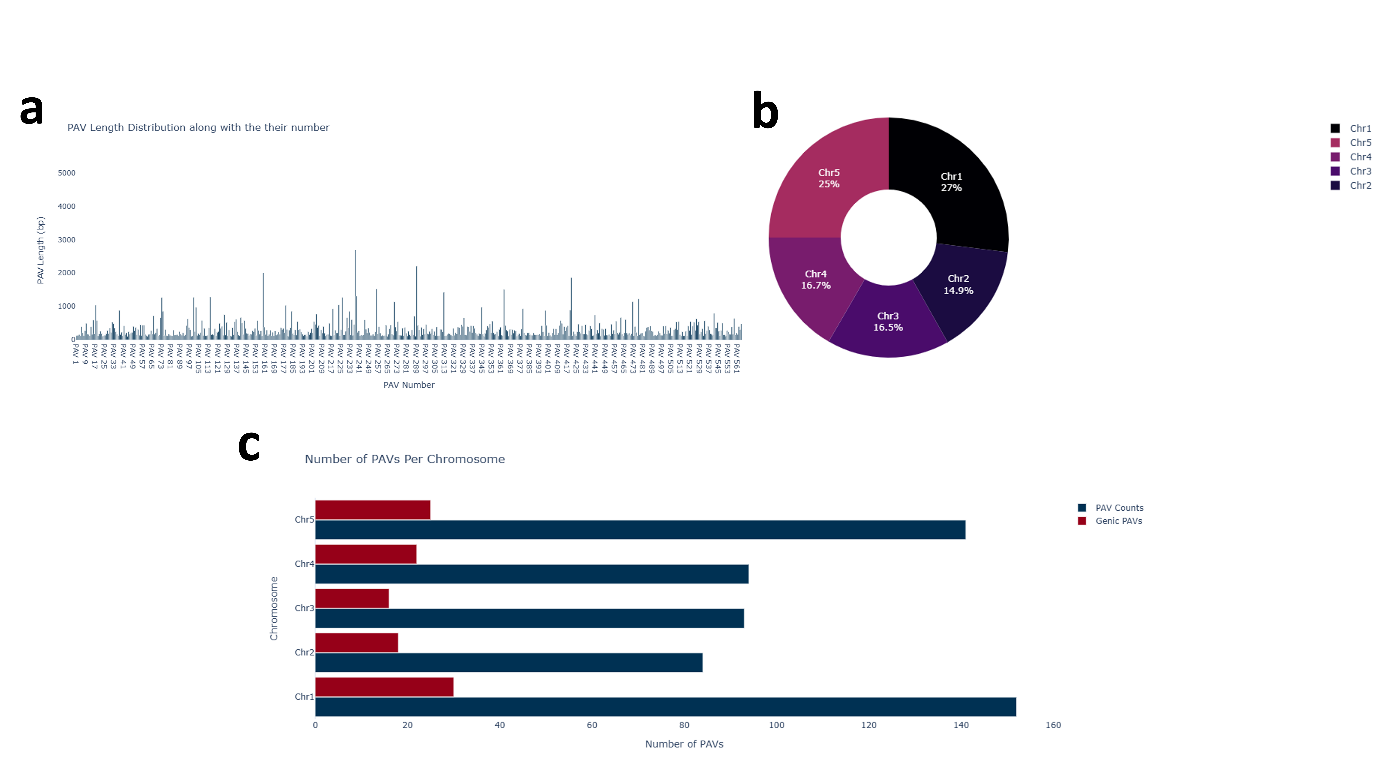

**Fig 3.10.** In the oy_0 assembly results, **(a)** PAV counts and lengths are presented, PAV number 237 is the longest variable region at 2.7 kb; **(b)** Chromosome 1 shows the highest variation, with its area covered by 152 PAVs, making it the most prominent variable region among all chromosomes. **(c)** PAVs are divided into genic and non-genic groups with counts.

**Ecotype_11_** **po_0**
Total PAVs: 727
Longest PAV: 2.3 kb

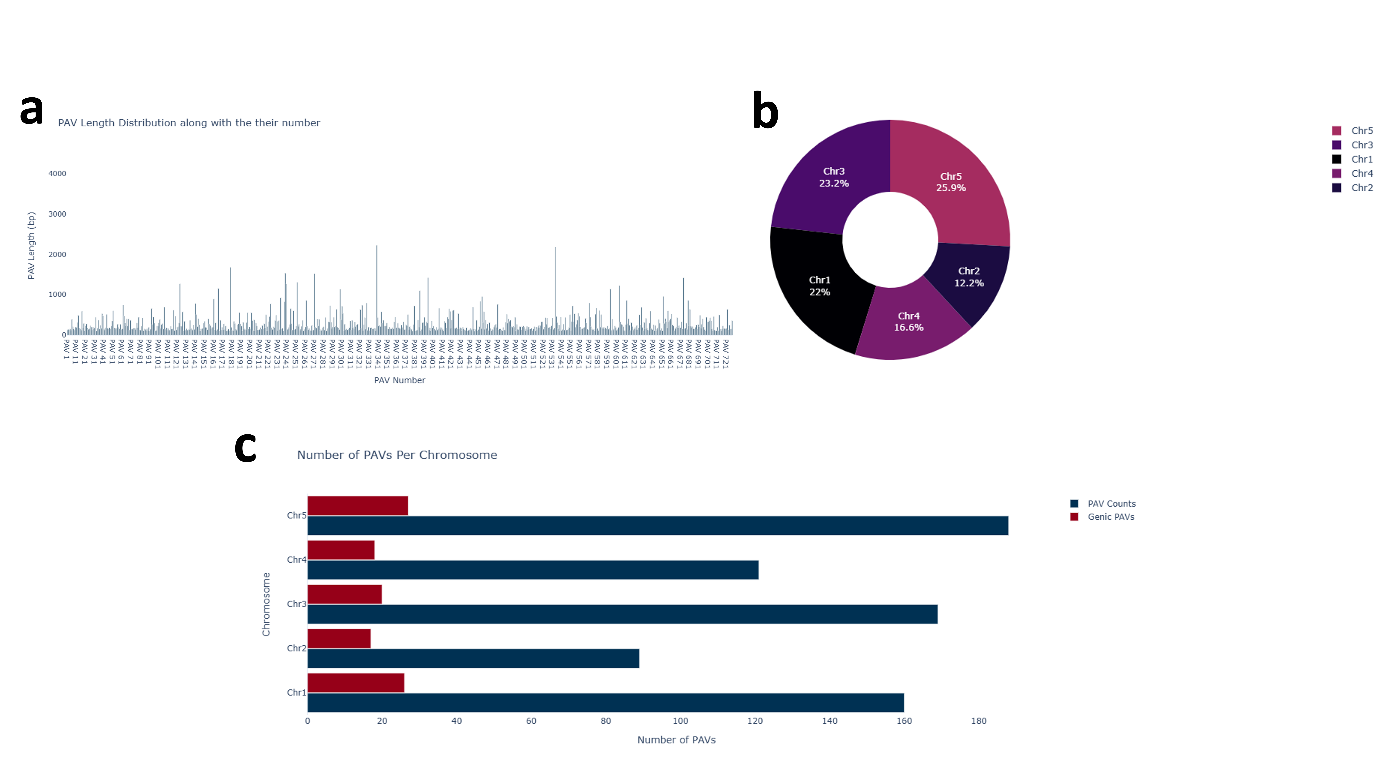

**Fig 3.11.** Result figures for the po_0 assembly: **(a)** Number of PAVs along with their lengths, highlighting both the shortest and longest detected regions—PAV number 339 is the longest chunk, though it originates from a chromosome 3 region; **(b)** Percentage of chromosomal area covered by PAVs; and **(c)** Classification of PAVs into genic and non-genic categories with their respective counts.

**Ecotype_12_rsch_4**
Total PAVs: 1070
Longest PAV: 3.8 kb

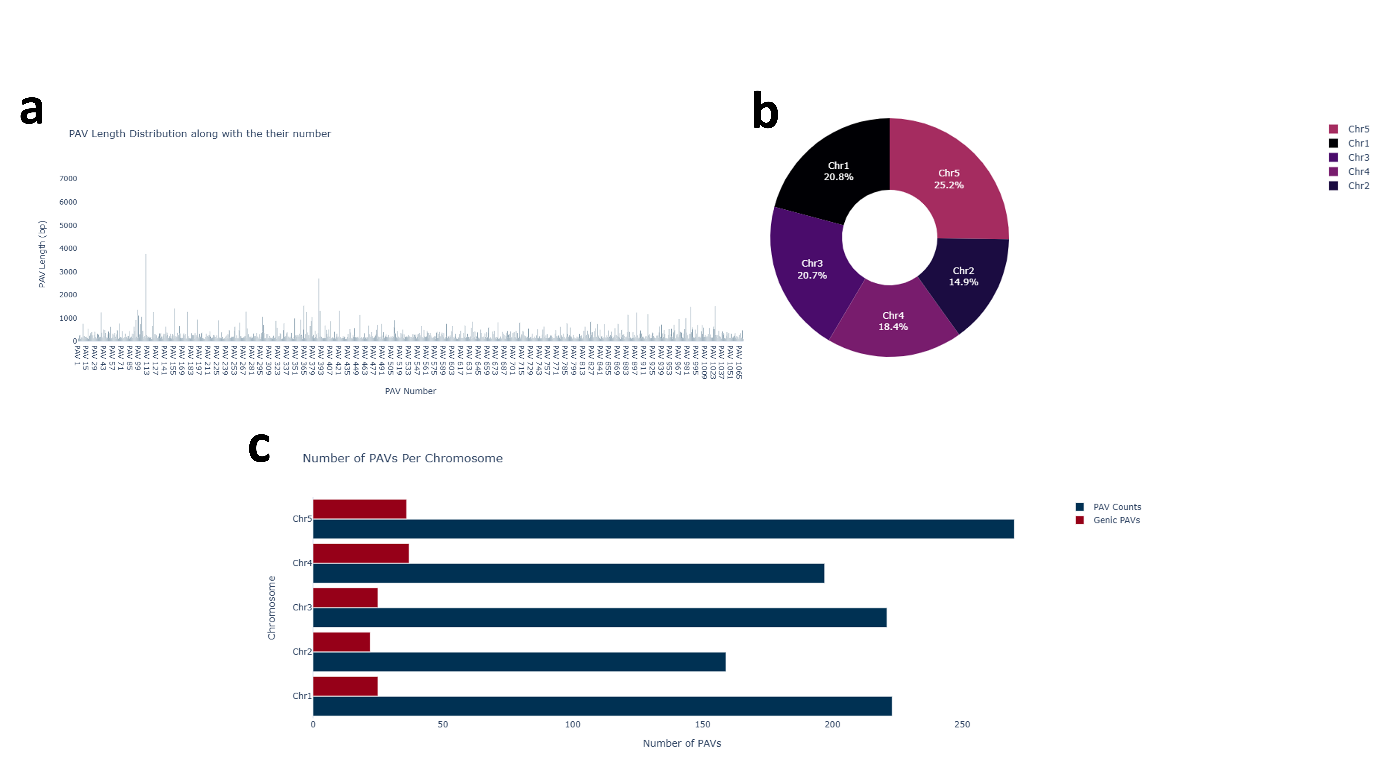

**Fig 3.12.** Result figures for the rsch_4 assembly: **(a)** Number of PAVs along with their lengths, highlighting both the shortest and longest detected regions; **(b)** Chromosomes 5 account for 25% of the total variation area covered by PAVs; and **(c)** Classification of PAVs into genic and non-genic categories with their respective counts. A total of 270 PAVs were detected on chromosome 5 in the rsch_4 assembly, representing the highest number of PAVs observed.

**Ecotype_13_sf_2**
Total PAVs: 1353
Longest PAV: 2.4 kb

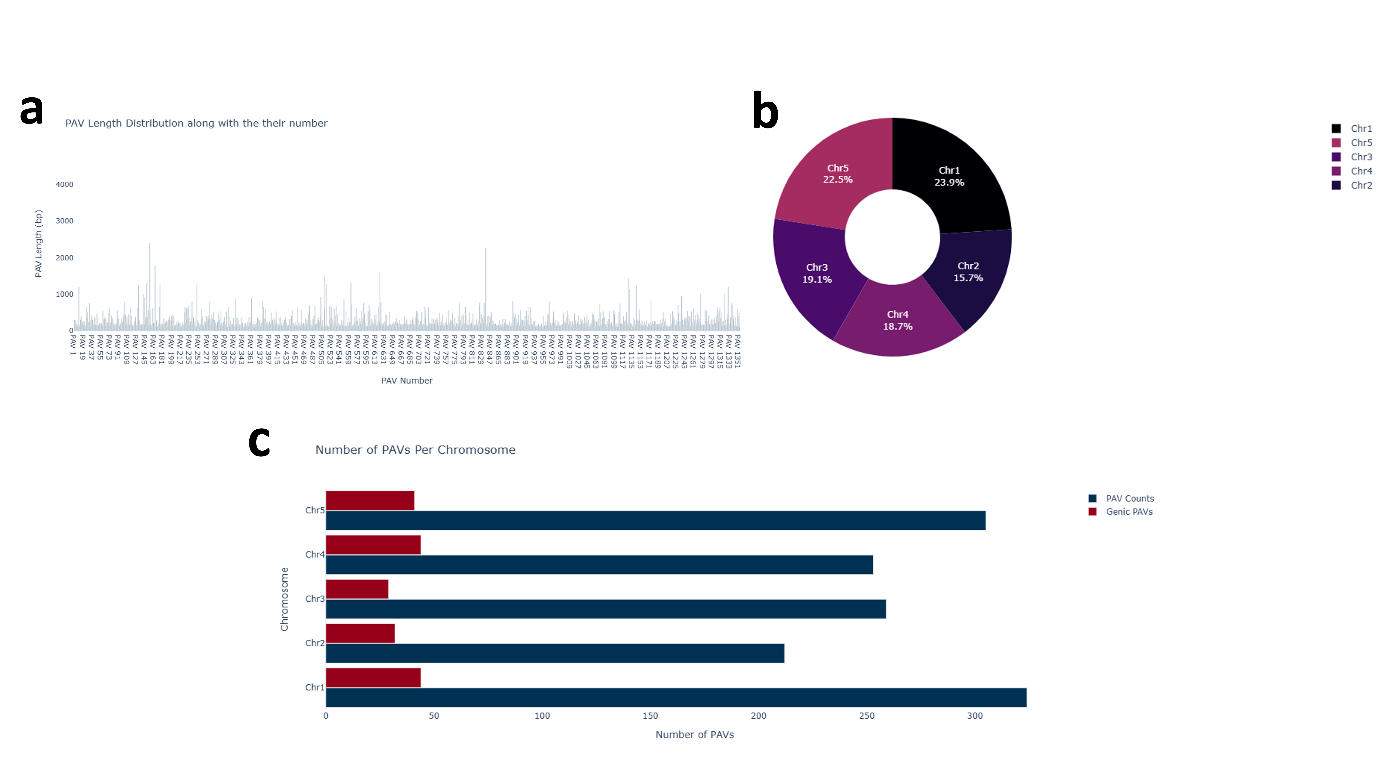

**Fig 3.13.** Result figures for the sf_2 assembly: **(a)** Number of PAVs along with their lengths, highlighting both the shortest and longest detected regions—PAV number 154 is the longest, measuring 2.4 kb; **(b)** Percentage of chromosomal area covered by PAVs; and **(c)** Classification of PAVs into genic and non-genic categories with their respective counts.

**Ecotype_14_tsu_0**
Total PAVs: 1163
Longest PAV: 2.7 kb

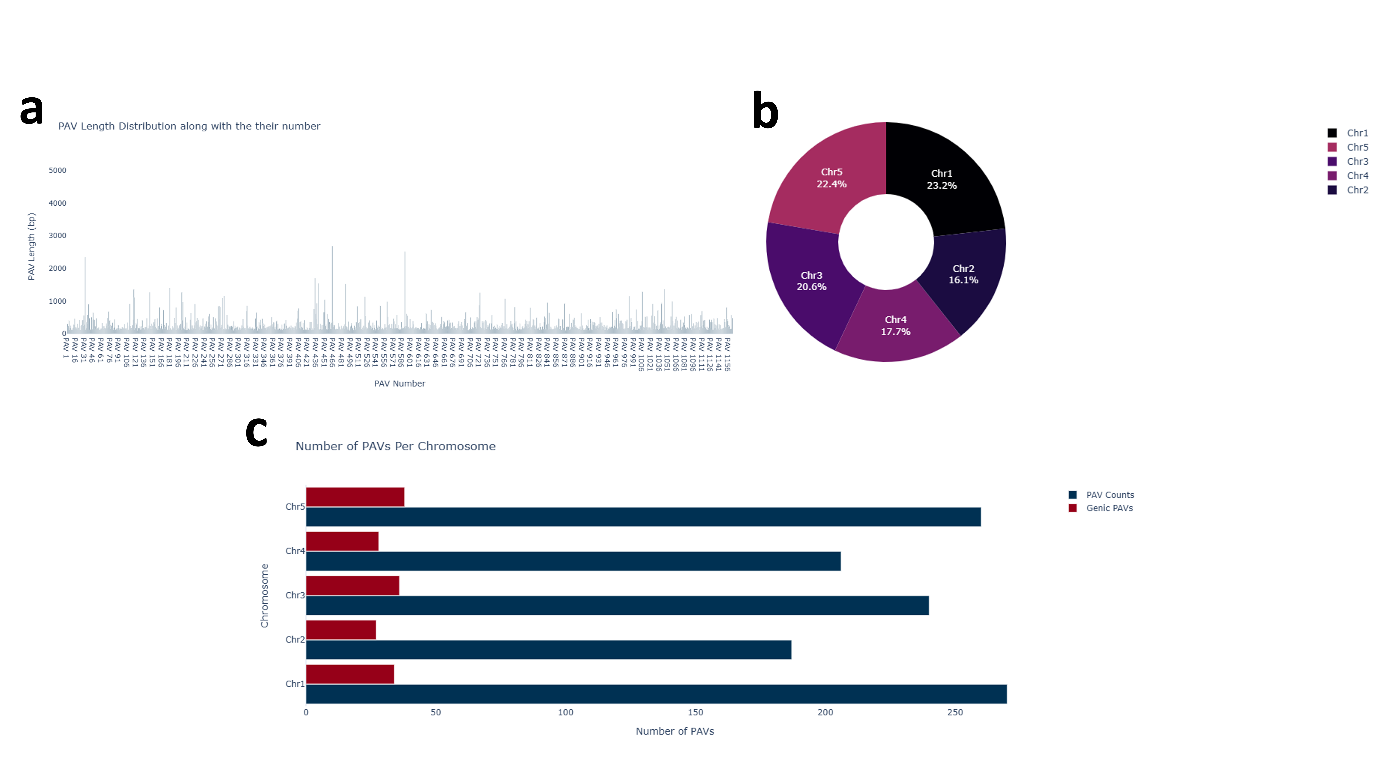

**Fig 3.14.** Result figures for the tsu_0 assembly: (a) Number of PAVs along with their lengths, highlighting both the shortest and longest detected regions—surprisingly, the PAV number 464 on chromosome 3 is the longest, measuring 2.7 kb; (b) Chromosome 2 exhibits the lowest variation, with only 16% of its area covered by PAVs; and (c) Classification of PAVs into genic and non-genic categories with their respective counts.

**Ecotype_15_wil_2**
Total PAVs: 1231
Longest PAV: 3.5 kb

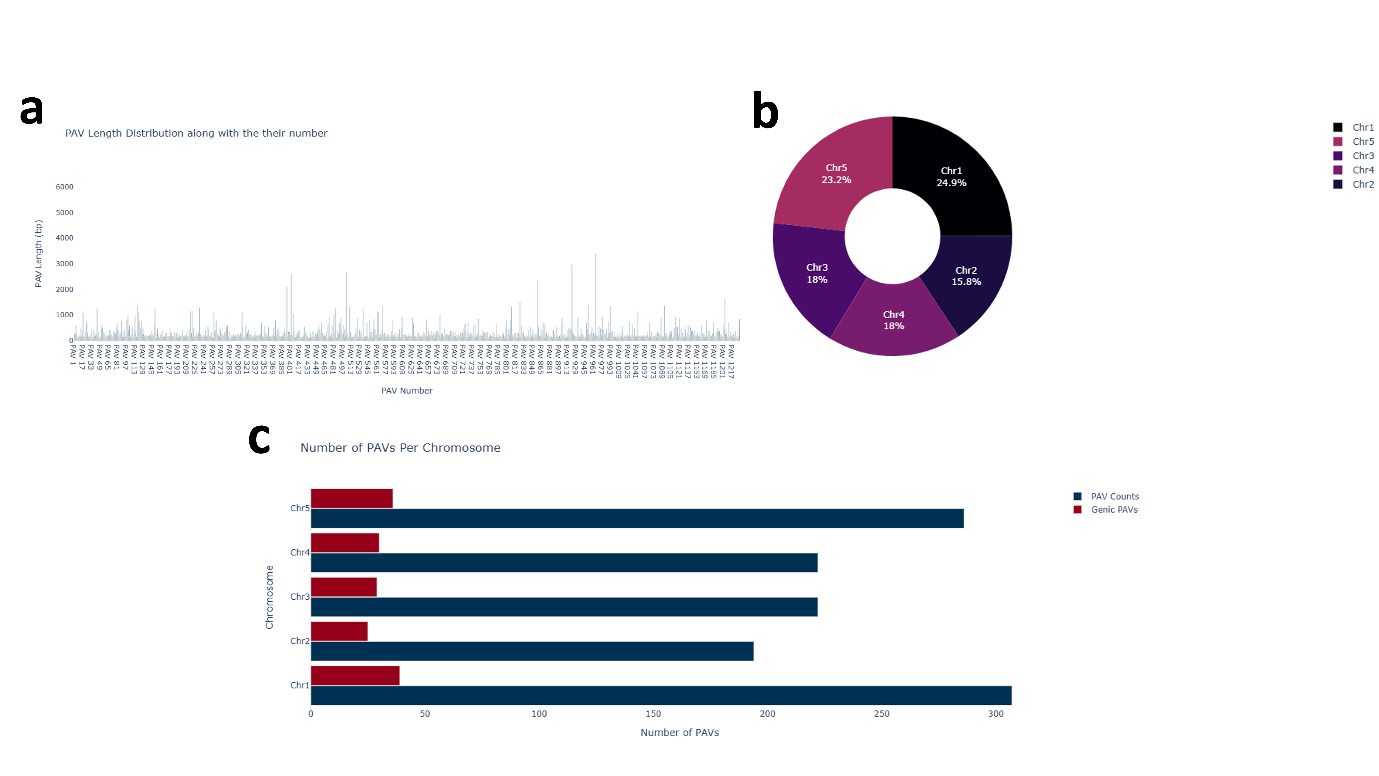

**Fig 3.15.** Result figures for the wil_2 assembly: (a) Distribution of PAV lengths across the genome, highlighting the shortest and longest regions—PAV number 965 is the longest, located in chromosome 5 region, measuring 3.5 kb; (b) Chromosomal distribution of PAVs, with chromosome 1 covering the largest area at 25% and chromosome 2 showing the lowest variation, covering 15 %; (c) Classification of PAVs into genic and non-genic categories, with 159 PAVs identified as genic, and 1072 PAVs identified as non-genic. Notably, 15 % of the PAVs overlap with coding regions, suggesting potential functional impacts.

**Ecotype_16_ws_0**
Total PAVs: 1155
Longest PAV: 2.7 kb

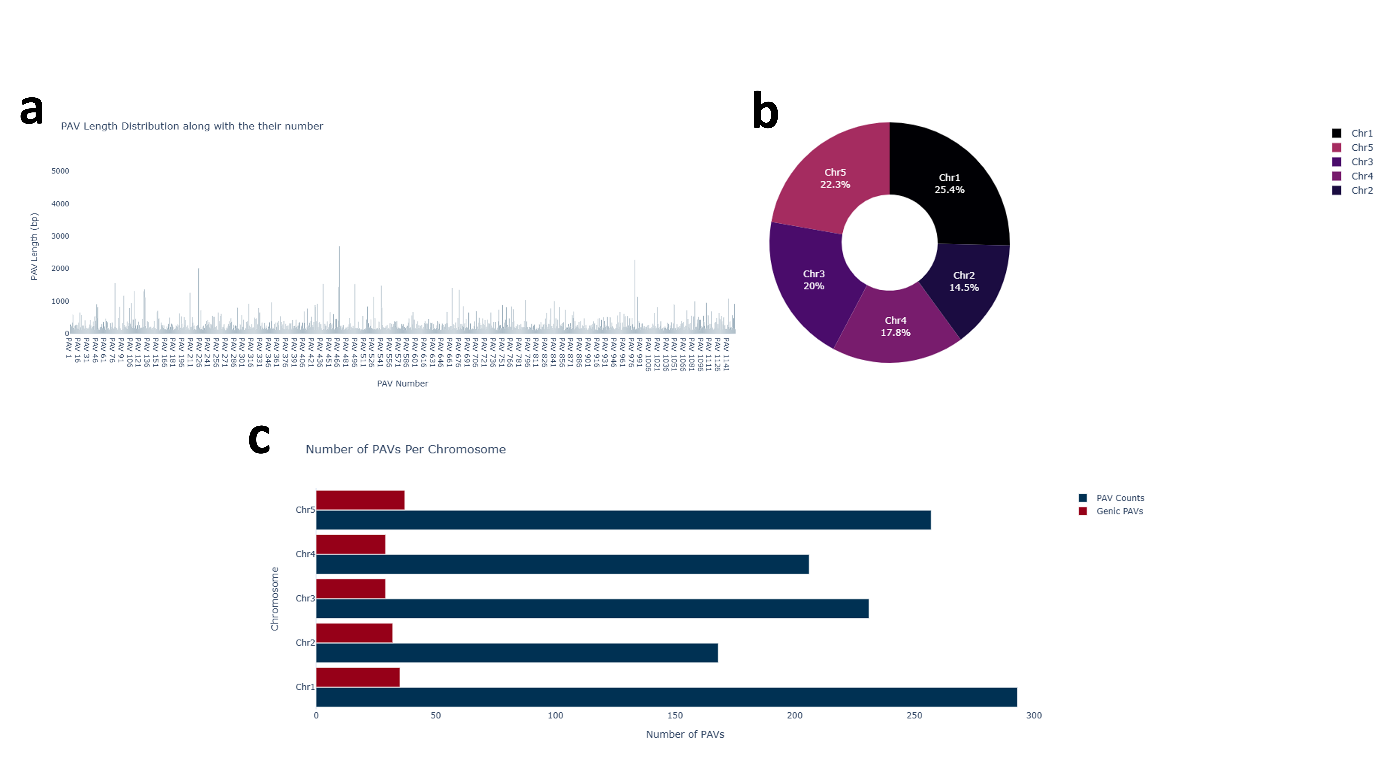

**Fig 3.16.** Result figures for the ws_0 assembly; **(a)** The number and lengths of PAVs are shown, with the shortest and longest regions highlighted. PAV number 468 is the longest, though it is located in chromosome 3 with a ~3 kb region; **(b)** The percentage of chromosomal area covered by PAVs is displayed; and **(c)** PAVs are classified into genic and non-genic categories, with their respective counts provided. A total of 293 PAVs were detected on chromosome 1 of the ws_0 assembly.

**Ecotype_17_wu_0**
Total PAVs: 935
Longest PAV: 2.7 kb

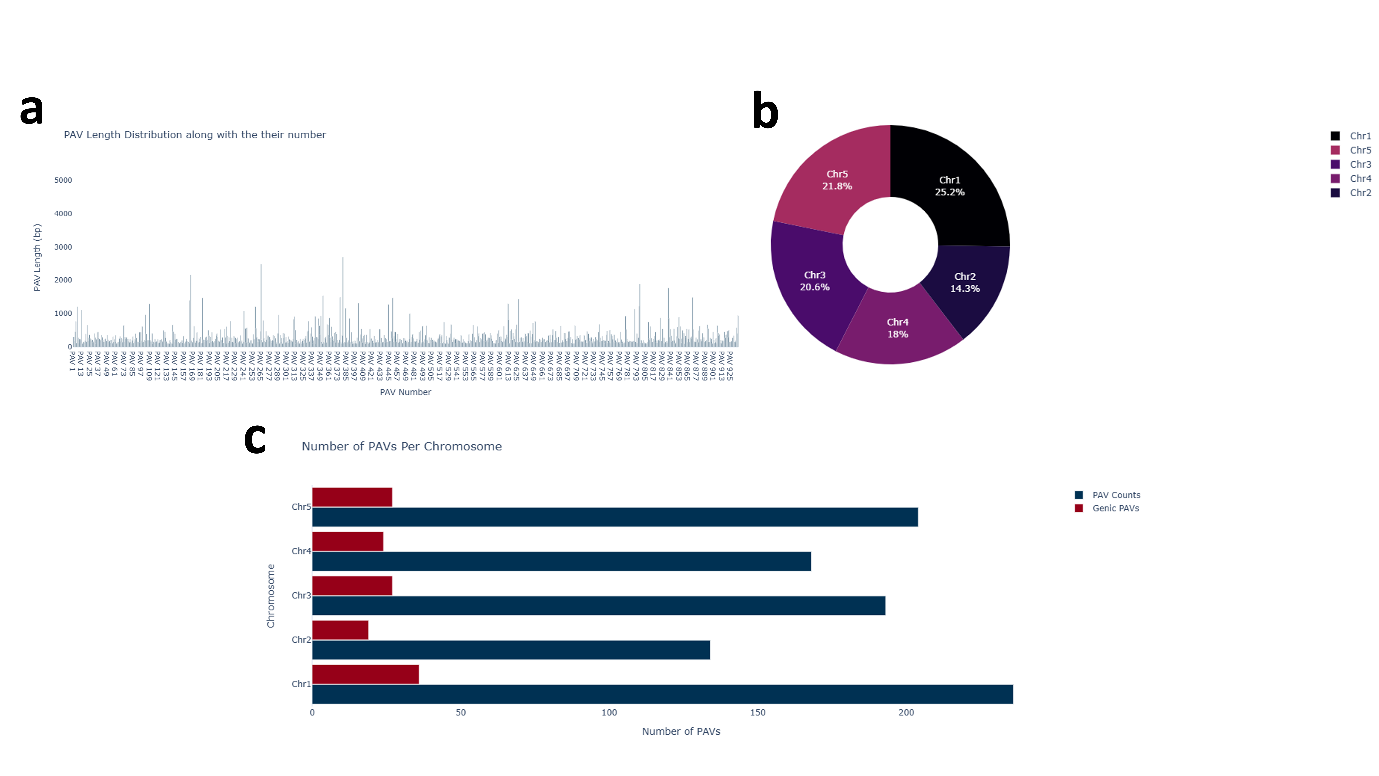

**Fig 3.17.** Result figures for the wu_0 assembly: **(a)** The number and lengths of PAVs are shown, highlighting both the shortest and longest detected regions; **(b)** The percentage of chromosomal area covered by PAVs; and **(c)** Classification of PAVs into genic and non-genic categories, with their respective counts in the wu_0 assembly.

**Ecotype_18_zu_0**
Total PAVs: 1068
Longest PAV: 3.7 kb

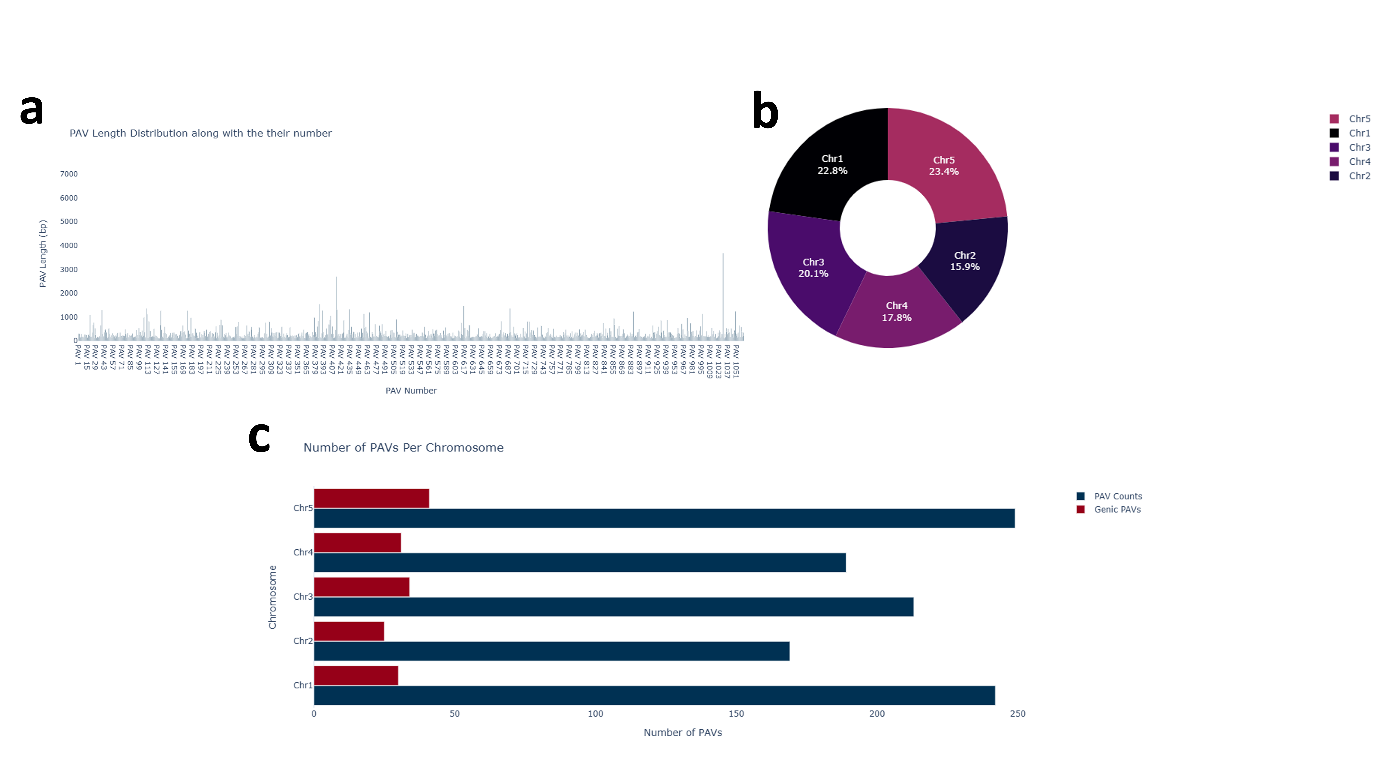

**Fig 3.18.** Result figures for the zu_0 assembly **(a)** Shows the number of PAVs with their length, highlighting both the shortest and longest detected regions, PAV number 1029 is longest with 3.7 kb chromosomal region **(b)** Percentage area covered by PAV on chromosomes **(c)** Classification of PAVs into genic and non-genic PAVs categories with their respective counts and 161 genic PAVs detected in zu_0 assembly covering 15% coding region as a variation.

3.2 *P. communis* (Pear) Cultivars PAV Analysis Summary

**Cultivar_1_Bartlett**
Total PAVs: 24,377
Average length of PAV: 5.7 kb

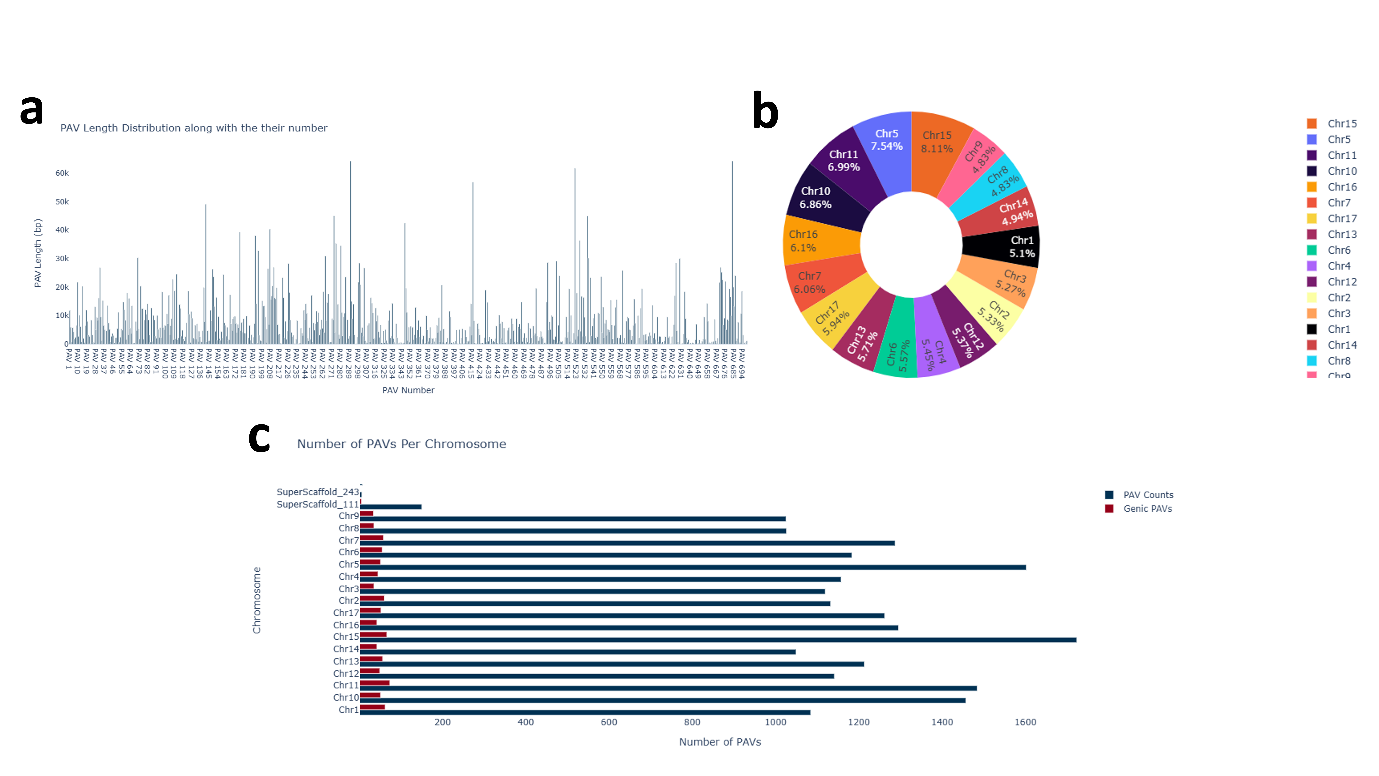

**Fig 3.19.** Result figures for the Bartlett_v2.0 genome **(a)** Shows the number of PAVs with their length, highlighting both the shortest and longest detected regions, **(b)** Percentage area covered by PAV on chromosomes **(c)** Classification of PAVs into genicPAVs and non-genicPAVs categories with their respective counts and 1602 PAVs detected on chr5 in Bartlett_v2.0.genome which are largest number of PAVs per chromosome.

**Cultivar_2_Cuiguan**
Total PAVs: 13,207
Average length of PAV: 7.1 kb

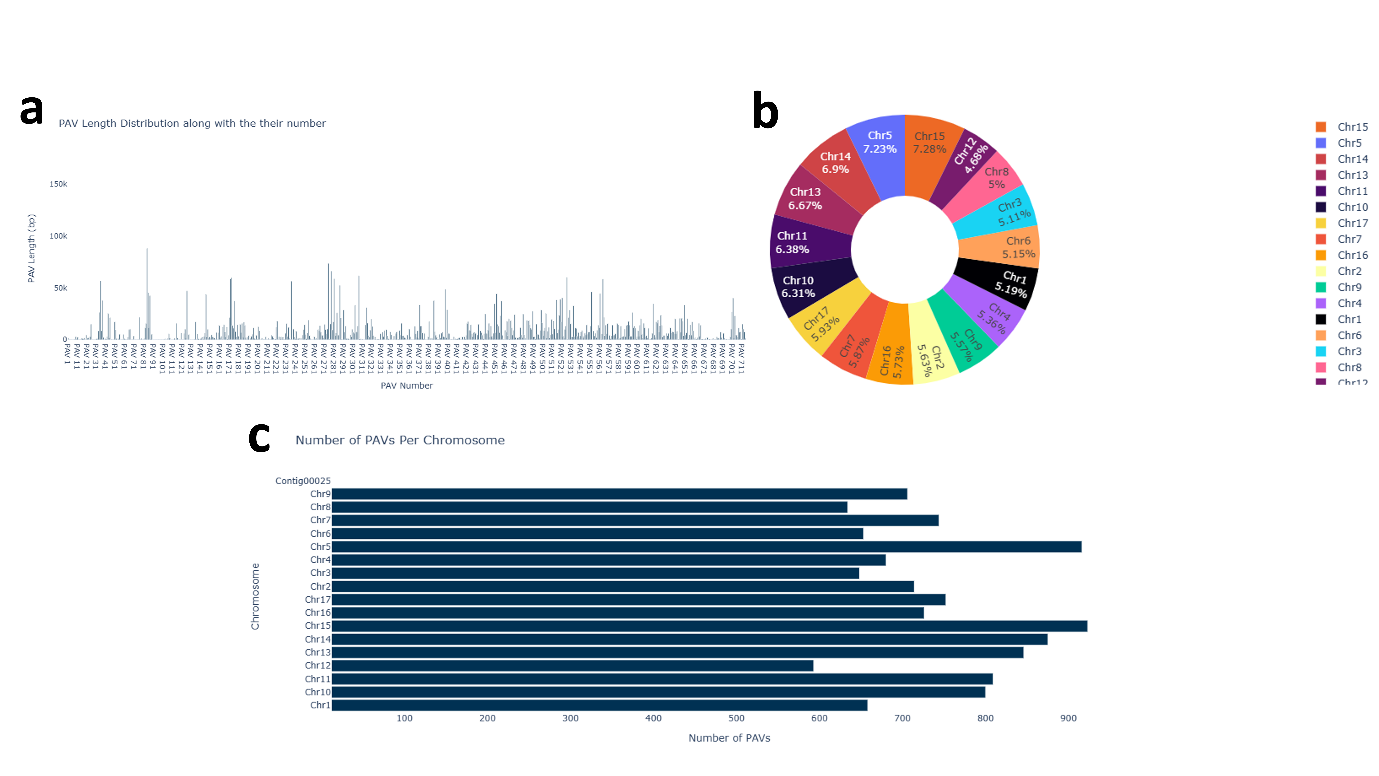

**Fig 3.20.** Result figures for the Cuiguan_v1.0 genome **(a)** Shows the number of PAVs with their length, highlighting both the shortest and longest detected regions, **(b)** 7.28 Percent area covered by PAV on chromosome no 15 which is the largest variation **(c)** Classification of PAVs into genic and non-genic PAVs categories with their respective counts

**Cultivar_3_Shanxiduli**
Total PAVs: 16,168
Average length of PAV: 5.4 kb

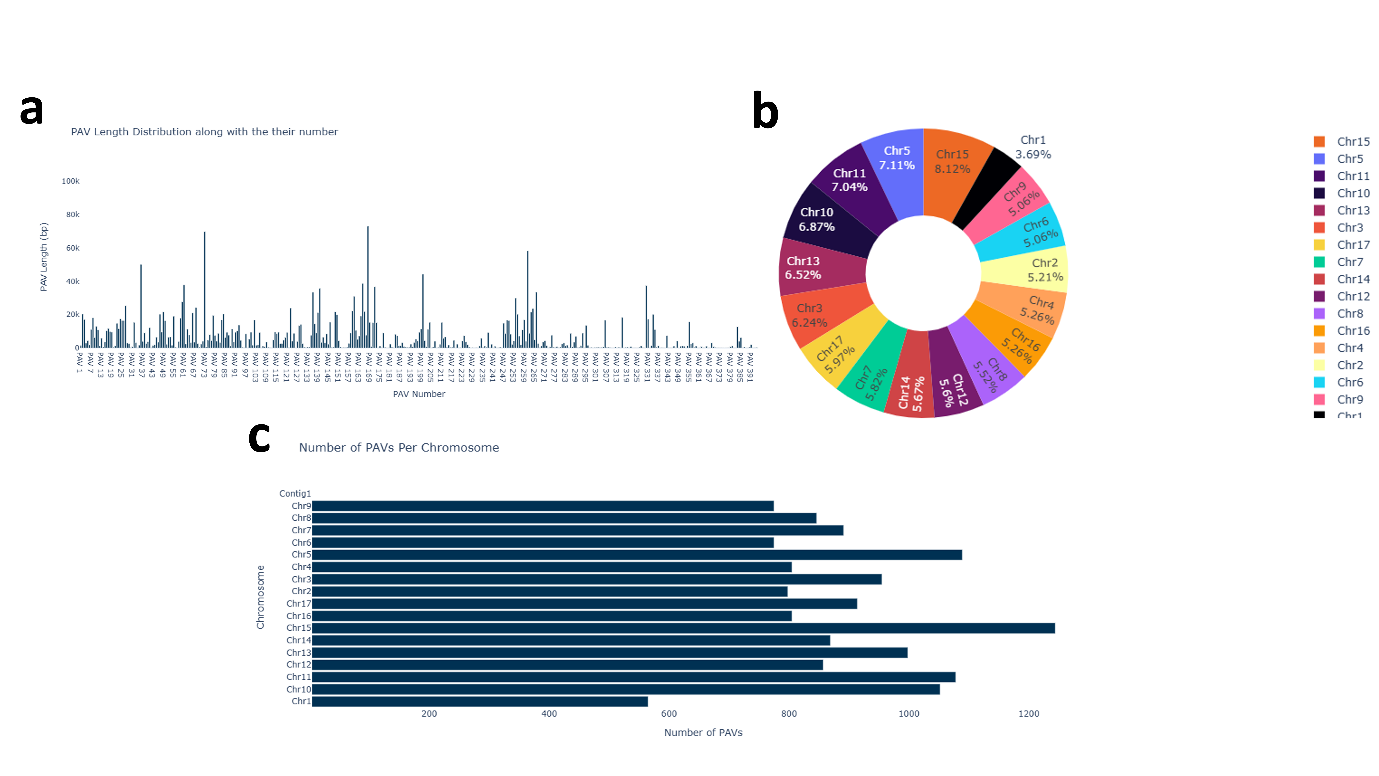

**Fig 3.21.** Result figures for the Shanxiduli genome **(a)** represent the number of PAVs with their length, highlighting both the shortest and longest detected regions **(b)** Percentage area covered by PAV on chromosomes **(c)** Classification of PAVs into genic and non-genic PAVs categories with their respective counts and notably there were no single genic PAV detected in Shanxiduli genome.

**Cultivar_4_ZhongaiNO.1**
Total PAVs: 16,720
Average length of PAV: 6.1 kb

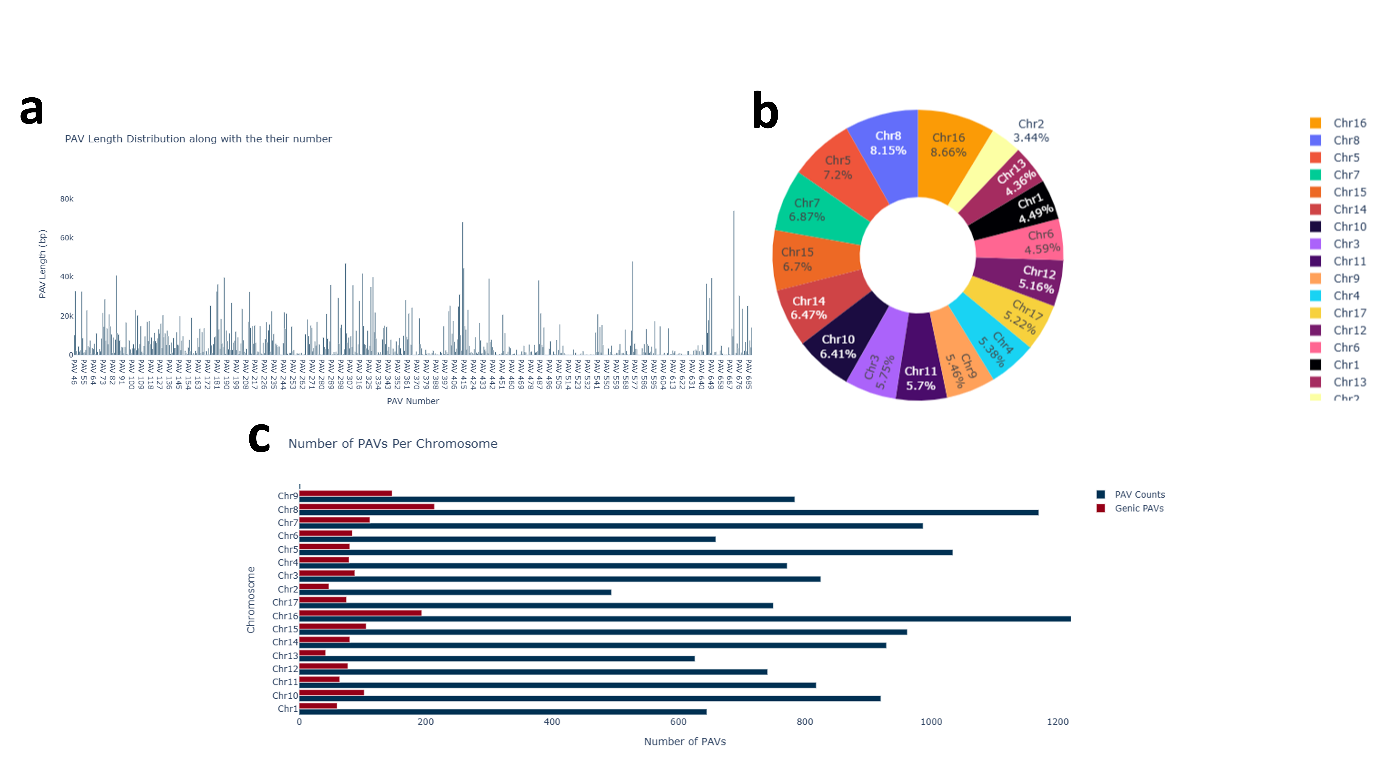

**Fig 3.22.** Result figures for the ZhongaiNO.1 genome **(a)** exhibit the number of PAVs with their length, highlighting both the shortest and longest detected regions. **(b)** Percentage area covered by PAV on chromosomes **(c)** Classification of PAVs into genic and non-genic PAV categories.

3.2 *M. musculus* (Mouse) PAV Analysis Summary

**Assembly _1_ 129S1_SvImJ**

Total PAVs: 4,824
Average length of PAV: 1.3 kb

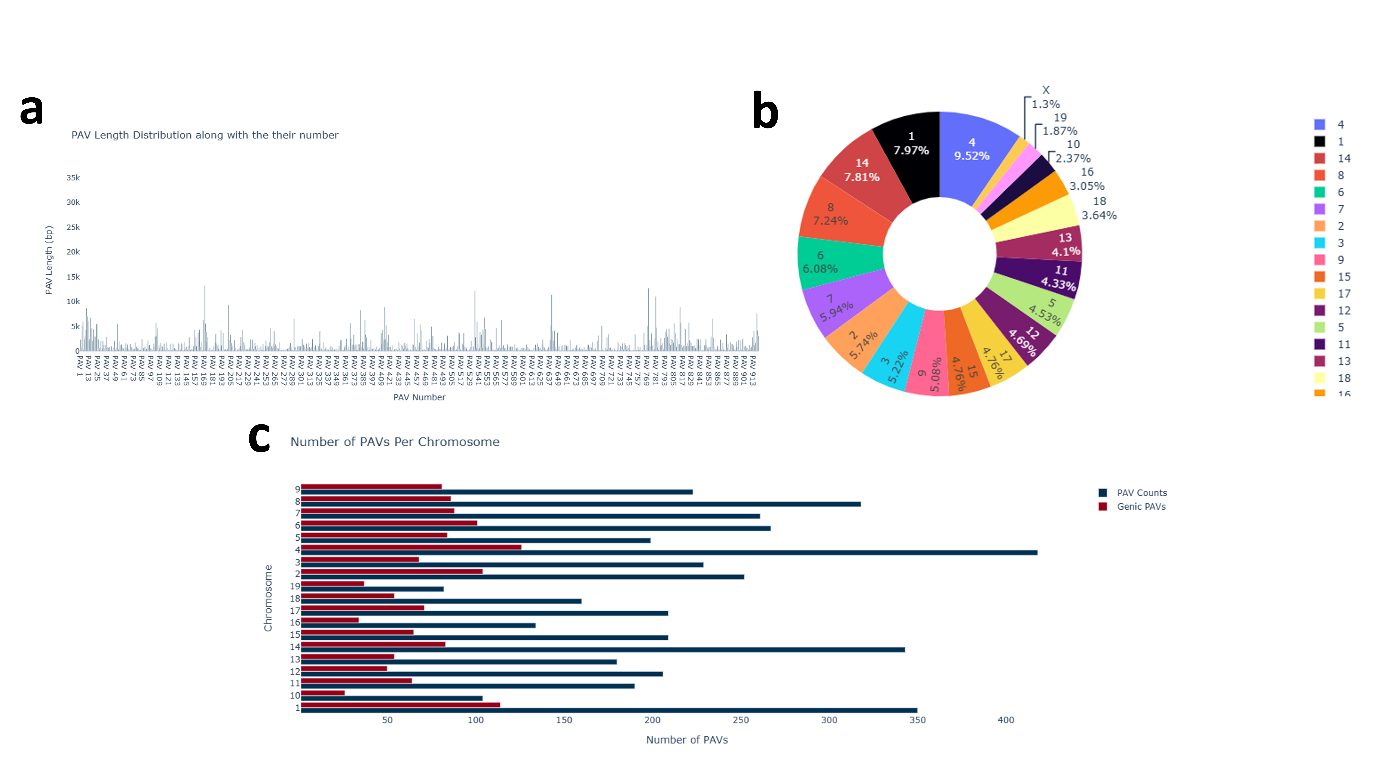

**Fig 3.23.** Result figures for the 129S1_SvImJ assembly **(a)** Shows the number of PAVs with their length, highlighting both the shortest and longest detected regions **(b)** Percentage area covered by PAV on chromosomes **(c)** Classification of PAVs into genic and non-genic PAVs categories with their respective counts and 1404 total PAVs detected which are involved in coding region 129S1_SvImJ assembly.

**Assembly_2_AKR_J**
Total PAVs: 5,080
Average length of PAV: 1.4 kb

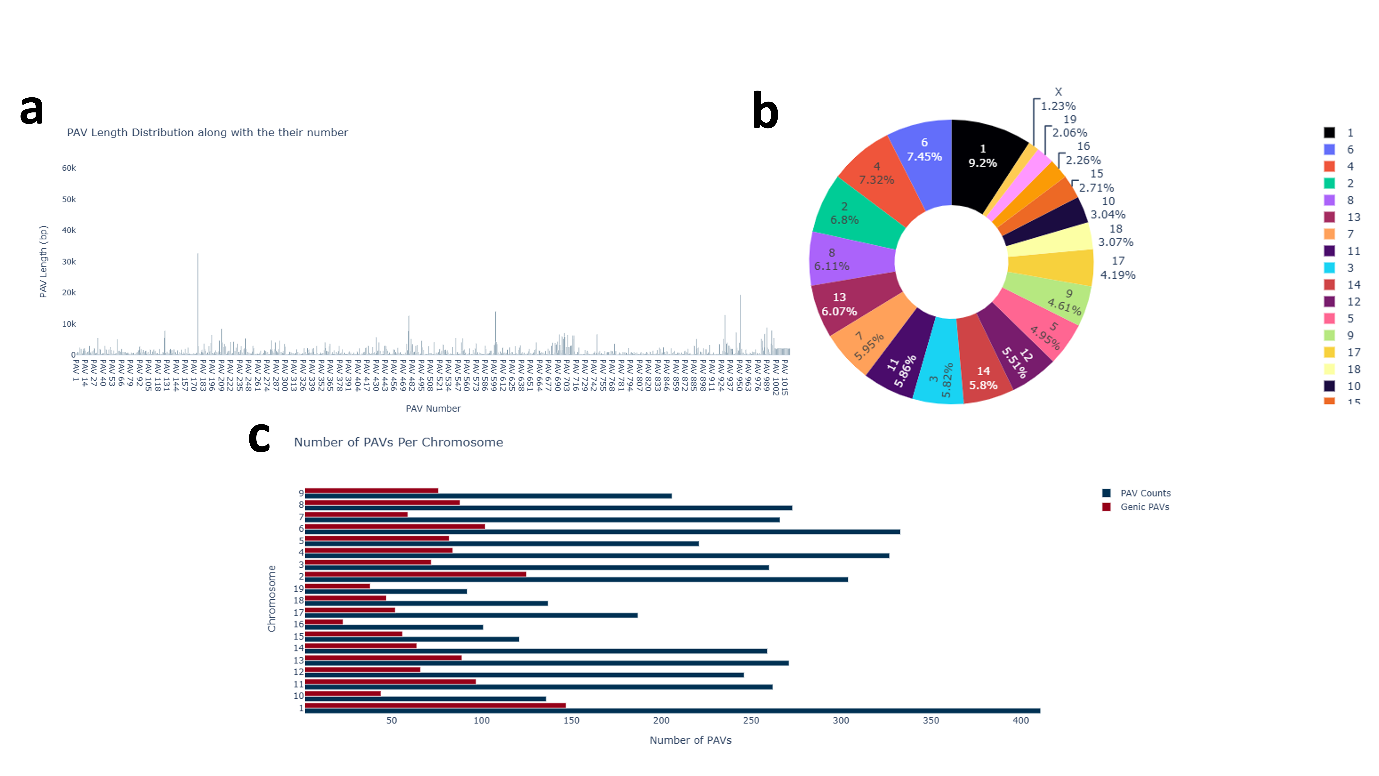

**Fig 3.24.** Result figures for the AKR_J assembly **(a)** show the number of PAVs with their length, highlighting both the shortest and longest detected regions; **(b)** Percentage area covered by PAV on chromosomes; **(c)** Classification of PAVs into genic PAVs and non-genic PAVs categories with their respective counts.

**Assembly _3_C3H_HeJ**
Total PAVs: 4,784
Average length of PAV: 1.2 kb

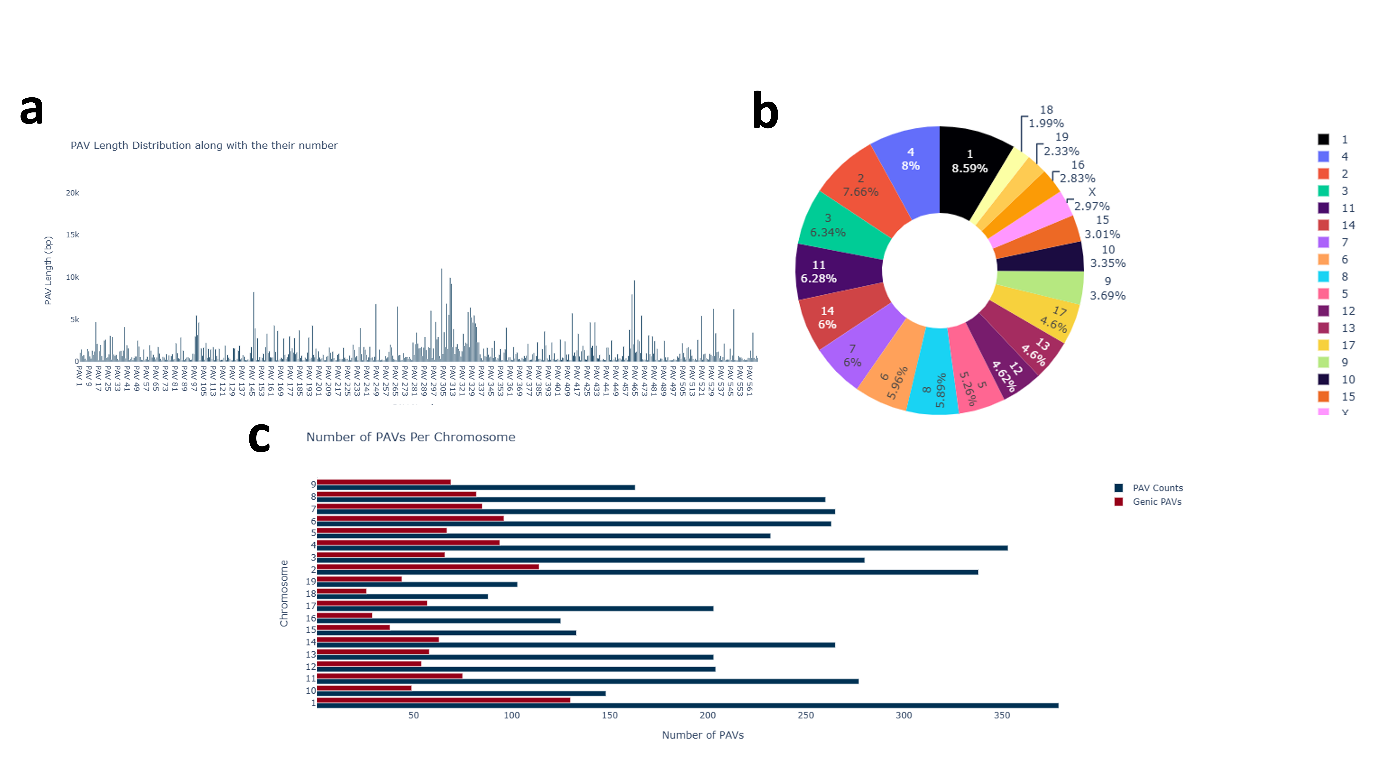

**Fig 3.25.** Result figures for the C3H_HeJ assembly **(a)** show the number of PAVs with their length, highlighting both the shortest and longest detected regions **(b)** 9 Percent area covered of PAVs by chromosome 1 alone **(c)** Classification of PAVs into genic PAVs and non-genic PAVs categories with their respective counts.

**Assembly _4_C57BL_6NJ**
Total PAVs: 749
Average length of PAV: 1.4 kb

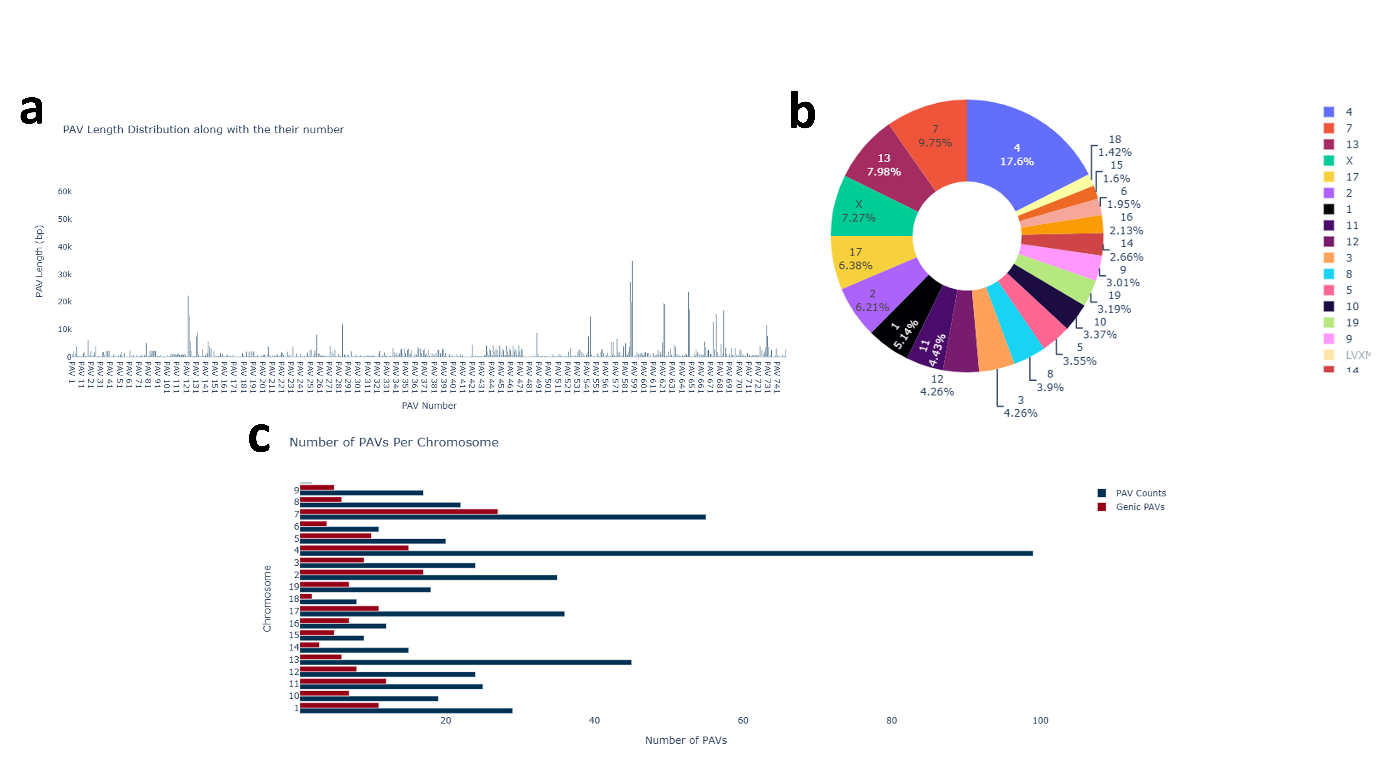

**Fig 3.26.** Result figures for the C57BL_6NJ assembly **(a)** Shows the number of PAVs with their length, highlighting both the shortest and longest detected regions, C57BL_6NJ_v1 has lowest number with 749 of PAVs as compared to other assemblies **(b)** Percentage area covered by PAV on chromosomes **(c)** Classification of PAVs into genicPAVs and non-genicPAVs categories with their respective counts. 175 PAVs intersect with genes in the C57BL_6NJ assembly.

**Assembly_5_DBA_2J**
Total PAVs: 5,156
Average length of PAV: 1.3 kb

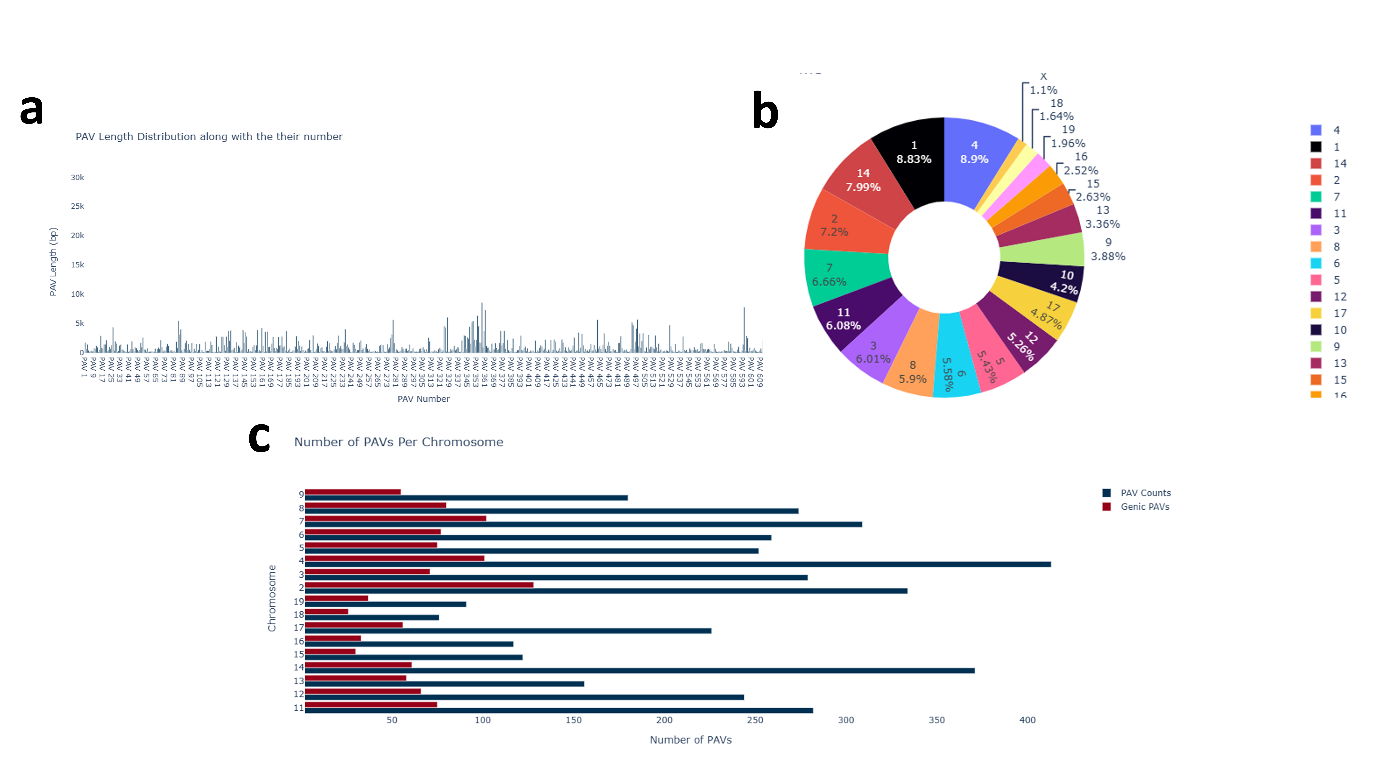

**Fig 3.27.** Result figures for the DBA_2J assembly **(a)** show the number of PAVs with their length, highlighting both the shortest and longest detected regions; **(b)** Percentage area covered by PAV on chromosomes; **(c)** Classification of PAVs into genic and non-genic PAVs categories with their respective counts.

**Table S1**. Presence-Absence Variation (PAV) numbers identified by XtractPAV in different organisms, concerning their assembly name and the count of PAVs.

| **Organism** | **Assembly Name** | **No. of PAVs** |
| --- | --- | --- |
| *M. musculus* | 129S1_SvImJ_v1 | 4,824 |
| *M. musculus* | AKR_J_v1 | 5,080 |
| *M. musculus* | C3H_HeJ_v1 | 4,784 |
| *M. musculus* | C57BL_6NJ_v1 | 749 |
| *M. musculus* | DBA_2J_v1 | 5,156 |
| *P. communis* (Pear) | Bartlett_v2.0 | 24,377 |
| *P. communis* (Pear) | Cuiguan_v1.0 | 13,207 |
| *P. communis* (Pear) | Shanxiduli | 16,168 |
| *P. communis* (Pear) | ZhongaiNO.1 | 16,720 |
| *A. thaliana* | bur_0.v7.fas | 1093 |
| *A. thaliana* | can_0.v7.fas | 1370 |
| *A. thaliana* | ct_1.v7.fas | 611 |
| *A. thaliana* | edi_0.v7.fas | 1201 |
| *A. thaliana* | hi_0.v7.fas | 450 |
| *A. thaliana* | kn_0.v7.fas | 1028 |
| *A. thaliana* | ler_0.v7.fas | 1015 |
| *A. thaliana* | mt_0.v7.fas | 543 |
| *A. thaliana* | no_0.v7.fas | 1038 |
| *A. thaliana* | oy_0.v7.fas | 564 |
| *A. thaliana* | po_0.v7.fas | 727 |
| *A. thaliana* | rsch_4.v7.fas | 1070 |
| *A. thaliana* | sf_2.v7.fas | 1353 |
| *A. thaliana* | tsu_0.v7.fas | 1163 |
| *A. thaliana* | wil_2.v7.fas | 1231 |
| *A. thaliana* | ws_0.v7.fas | 1155 |
| *A. thaliana* | wu_0.v7.fas | 295 |
| *A. thaliana* | zu_0.v7.fas | 1062 |

3.2 *Salmonella Enterica* serovars PAV Analysis

We have conducted the prokaryotic analysis with the *S. enterica* serovars. We used all *S. enterica* serovars genomes from the study (Jacobsen, et al., 2011) and S. enterica Typhimurium *str. LT2* as a reference, and the rest of the genomes were used as the query. The result of each serovar is shown in Table S2 below.

**Table S2.** PAV Counts across 41 Salmonella Enterica Serovars

| **Organism** | **Accession** | **PAVs** |
| --- | --- | --- |
| *S. arizonae 62:z4,z23* | GCA_000018625.1 | 308 |
| *S. Agona SL483* | GCA_000020885.1 | 107 |
| *S. Dublin CT_02021853* | GCA_000020925.1 | 78 |
| *S. Typhimurium D23580* | GCA_000027025.1 | 7 |
| *S. Kentucky CVM29188* | GCA_000170195.2 | 119 |
| *S. Saintpaul SARA23* | GCA_000170215.1 | 23 |
| *S. Saintpaul SARA29* | GCA_000170235.1 | 103 |
| *S. 4,[5],12:i:- CVM23701* | GCA_000170255.1 | 11 |
| *S. Javiana GA_MM04042433* | GCA_000171255.1 | 124 |
| *S. Kentucky CDC 191* | GCA_000171275.1 | 102 |
| *S. Schwarzengrund SL480* | GCA_000171295.1 | 121 |
| *S. Heidelberg SL486* | GCA_000171315.1 | 48 |
| *S. Newport SL317* | GCA_000171415.1 | 66 |
| *S. Hadar RI_05P066* | GCA_000171515.1 | 65 |
| *S. Virchow SL491* | GCA_000171535.2 | 82 |
| *S. Typhi E00-7866* | GCA_000180215.1 | 144 |
| *S. Typhi E01-6750* | GCA_000180235.1 | 138 |
| *S. Typhi E02-1180* | GCA_000180255.1 | 131 |
| *S. Typhi E98-0664* | GCA_000180275.1 | 143 |
| *S. Typhi E98-2068* | GCA_000180295.1 | 153 |
| *S. Typhi J185* | GCA_000180315.1 | 141 |
| *S. Typhi M223* | GCA_000180335.1 | 154 |
| *S. Typhi E98-3139* | GCA_000180375.1 | 127 |
| *S. Typhi CT18* | GCA_000195995.1 | 124 |
| *S. Typhimurium SL1344* | GCA_000210855.2 | 11 |
| *S. Tennessee CDC07-0191* | GCA_000353585.1 | 111 |
| *S. Typhimurium DT104* | GCA_001350055.1 | 09 |
| *S. Typhimurium 14028S* | GCA_008244785.1 | 4 |
| *S. Typhi 404ty* | GCA_010006625.1 | 106 |
| *S. Typhi AG3* | GCA_024028975.1 | 52 |
| *S. Typhi Ty2* | GCF_000007545.1 | 117 |
| *S. Choleraesuis SC-B67* | GCF_000008105.1 | 87 |
| *S. Enteritidis P125109* | GCF_000009505.1 | 68 |
| *S. Gallinarum 287/91* | GCF_000009525.1 | 56 |
| *S. Paratyphi A ATCC 9150* | GCF_000011885.1 | 101 |
| *S. Newport SL254* | GCF_000016045.1 | 66 |
| *S. Paratyphi C RKS4594* | GCF_000018385.1 | 73 |
| *S. Paratyphi B SPB7* | GCF_000018705.1 | 74 |
| *S. Heidelberg SL476* | GCF_000020705.1 | 61 |
| *S. Schwarzengrund CVM19633* | GCF_000020745.1 | 120 |
| *S. Paratyphi A AKU_12601* | GCF_000026565.1 | 103 |

**4. Data Availability**

XtractPAV is available at [Github_repository](https://github.com/SherazAhmadd/XtractPAV). The webpage of XtractPAV is [XtractPAV](https://sherazahmadd.github.io/XtractPAV/).
